# A divergent kinase lacking the glycine-rich loop regulates membrane ultrastructure of the *Toxoplasma* parasitophorous vacuole

**DOI:** 10.1101/397489

**Authors:** Tsebaot Beraki, Hu Xiaoyu, Malgorzata Broncel, Joanna C. Young, William J. O’Shaughnessy, Dominika M. Borek, Moritz Treeck, Michael L. Reese

## Abstract

Apicomplexan parasites replicate within a protective organelle called the parasitophorous vacuole (PV). The *Toxoplasma gondii* PV is filled with a network of tubulated membranes, which are thought to facilitate trafficking of effectors and nutrients. Despite being critical to parasite virulence, there is scant mechanistic understanding of the network’s functions. Here, we identify the parasite secreted kinase WNG1 as a critical regulator of tubular membrane biogenesis. WNG1 family members adopt an atypical protein kinase fold lacking the glycine rich ATP-binding loop that is required for catalysis in canonical kinases. Unexpectedly, we find that WNG1 is an active protein kinase that localizes to the PV lumen and phosphorylates PV-resident proteins, several of which are essential for the formation of a functional intravacuolar network. Moreover, we show that WNG1-dependent phosphorylation of these proteins is required for their membrane association, and thus their ability to tubulate membranes. Consequently, WNG1 knockout parasites have an aberrant PV membrane ultrastructure. Collectively, our results describe a unique family of *Toxoplasma* kinases and implicate phosphorylation of secreted proteins as a mechanism of regulating PV formation during parasite infection.

## Introduction

Protein phosphorylation is the most common post-translational modification in eukaryotic cells. The addition and removal of specific phosphates is a key mediator of cellular information processing and signal transduction. Phosphorylation is catalyzed by protein kinases, which form one of the largest families of enzymes in mammals (1). The interface between an intracellular pathogen and its host cell is a special case in cellular signaling that defines both a pathogen’s ability to manipulate its host and the host’s ability to respond to and control the pathogen. The parasite *Toxoplasma gondii* is one of the most successful pathogens in the world, as it can infect virtually any cell type of almost all warm-blooded animals, including approximately one third of humans worldwide (2). *Toxoplasma* directly manipulates signaling at the host-pathogen interface by secreting a variety of effector proteins (3, 4), including ~50 protein kinases and pseudokinases (5, 6). However, the functions of most of these effectors are unknown.

One vital role for these secreted kinases is to maintain the parasite’s replicative niche within its host cell. Like many intracellular pathogens, *Toxoplasma* survives in a specialized membranous organelle called the parasitophorous vacuole (PV). This vacuole is maintained as distinct from host endosomal trafficking, and is protected from fusion with host lysosomes (7). Disruption of the PV membrane by host immune defenses leads to parasite death (8, 9), and the parasite has evolved effector molecules that can protect it from such host attacks (10, 11). Far from being an impermeable wall, however, the parasite selectively exports (12) and imports (13, 14) molecules across the PV membrane.

One of the most striking features of the PV is the intravacuolar network (IVN) of membranous tubules of 20-50 nm diameter that appear to bud from the PV membrane into the vacuolar lumen (15). Notably, the inside of the tubules is topologically contiguous with the host cytosol (15). The IVN has been associated with diverse phenomena, including nutrient uptake via trafficking of host-derived vesicles (16, 17), “ingestion” of soluble host proteins by the parasite (18), protection from antigen presentation (19), and a means by which parasite effectors localize to the PV membrane (20) and thus protect it destruction by host immune effectors (21). The dense granular proteins GRA2 or GRA6 are required for IVN biogenesis and parasites that lack either protein grow in vacuoles without the well-structured membranous tubules. While IVN-deficient parasites grow normally in *in vitro* cell culture, they have strongly attenuated virulence in a mouse model of infection (22).

The PV is thus a complex cellular compartment that mediates sophisticated, multidirectional trafficking, though the molecules that regulate its functions are largely a mystery. Many of the known components of the PV, and of the IVN in particular, are highly phosphorylated after they have been secreted from the parasite (23). About one third of the *Toxoplasma* kinome contains signal peptides but lack transmembrane domains, and are thus predicted to be secreted. Most of these kinases belong to a parasite-specific family that includes a number of virulence effectors (24, 25, 10) secreted into the host cytosol from the parasite rhoptries during invasion (26), and have been dubbed the “rhoptry kinase (ROPK)” family. A previous bioinformatic effort annotated the majority of predicted secreted kinases in *Toxoplasma* as ROPKs (5). Notably, vertebrate or ROPK effector kinases localized in the host cytosol cannot access PV-resident proteins on the lumenal side of the PV membrane. However, two members of the ROPK family, ROP21/27, were recently found to localize to a different set of secretory organelles, the dense granules, which secrete into the PV lumen. Because ROP21/27 are expressed mainly during the chronic stage of the parasite (27), they are unlikely to function in the regulation of processes during the acute stage, such as the biogenesis of the IVN.

In the present work, we identify a specialized family of kinases that lack the glycine-rich loop that is critical for nucleotide-binding in canonical kinases, leading us to name them the With-no-Gly-loop, or WNG, family. These WNG kinases are conserved throughout the coccidian family of parasites to which *Toxoplasma* belongs and are secreted into the PV. We solved the crystal structure of a family member which demonstrates that the N-lobe of the kinase does indeed lack the structural elements that form the Gly-loop. We found that at least one member of the family, WNG1/ROP35, is catalytically active and we identified a number of proteins associated with the IVN membrane as phosphorylated in a WNG1-dependent manner. Finally, we demonstrated that loss of these phosphorylation sites correlates with aberrant PV ultrastructure, likely due to the loss membrane association of proteins that drive the biogenesis of IVN tubules. Taken together, our data show the WNG family of kinases mediates specialized functions in regulating the proteins that create and maintain the coccidian host-parasite vacuolar interface.

## Results

### Identification of a divergent family of coccidian secreted kinases that lack the canonical glycine-rich loop

We reasoned that regulatory phosphorylation of PV-resident proteins would most likely be carried out by a conserved resident protein kinase that is secreted from the parasites dense granules. To identify potential PV-resident kinases, we compared the sequences of the predicted secreted kinases in *Toxoplasma*. We were surprised to find that a small family of parasite kinases appear to completely lack the glycine-rich, or P-loop, that is found in all canonical kinases and is required for binding the ATP in the active site (28, 29) (Supplemental Fig S1a,b). These kinases include three proteins annotated as ROPKs (ROP33, ROP34, and ROP35), and a pseudokinase, BPK1, that has previously been identified as PV resident and a component of the bradyzoite cyst wall (30). Phylogenetic analysis gave clear support for these proteins forming a clade that is distinct from canonical protein kinases (Figure 1), including the parasite ROPKs. Furthermore, we identified members of this family in every species of coccidian parasite for which genomic sequence is available (Figure 1 and Supplemental Table S1c), suggesting that they play an important role in the parasites’ pathogenic lifestyle. Notably, the majority of ROPKs are not conserved throughout coccidian parasites, with the exception of the PV-resident kinases ROP21/27 (31). Given the lack of the glycine-rich loop and phylogenetic evidence that indicates that these proteins form a distinct clade, we propose that the family be named the WNG (With-No-Gly-loop) kinases.

**Figure 1:**
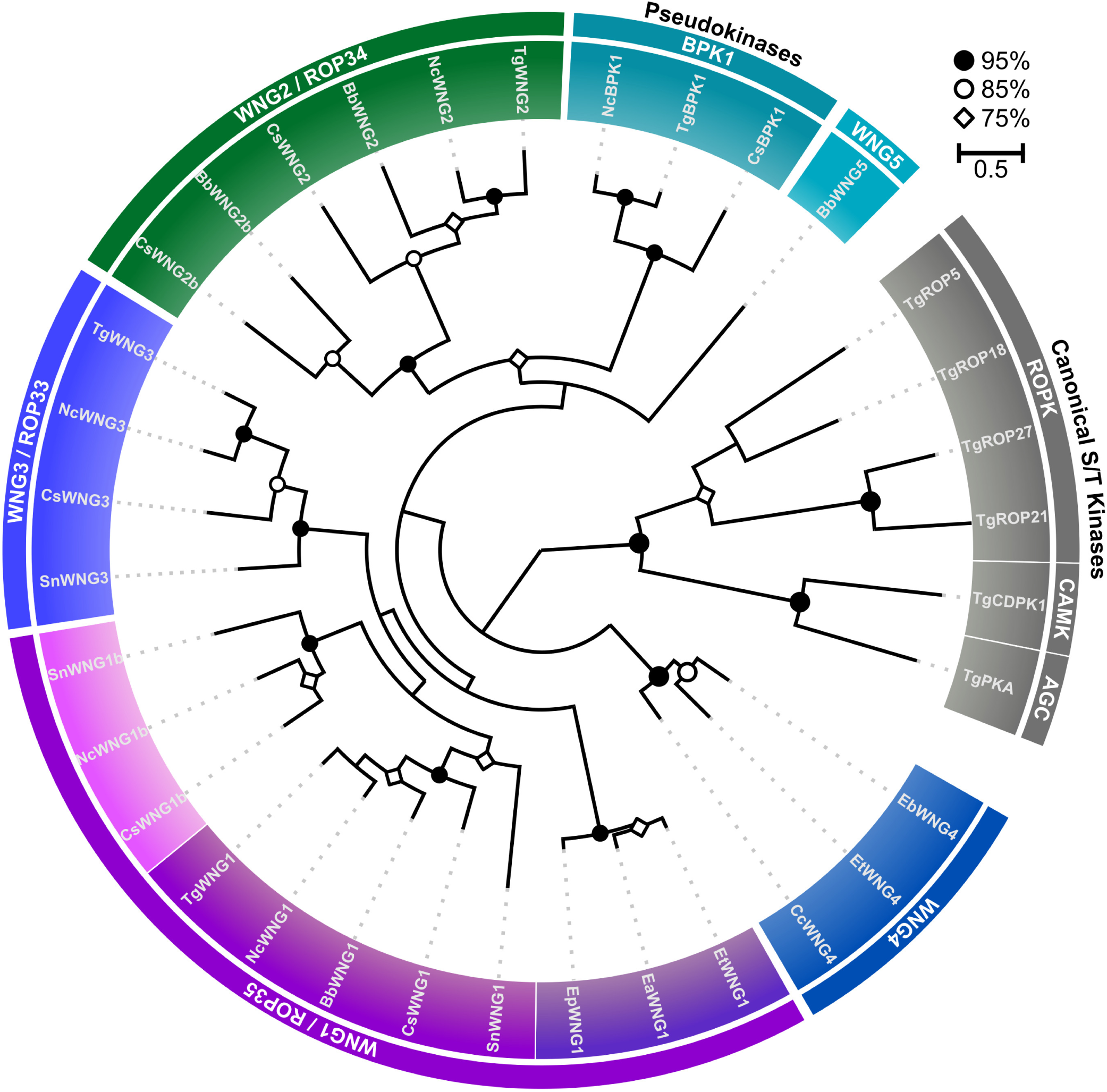
The WNG kinases comprise a phylogenetic clade that is distinct from canonical protein kinases. A maximum-likelihood phylogenetic tree estimated from the multiple sequence alignment of the indicated kinases. Bootstrap values are indicated as black circles (>95%); white circles (>85%); and white diamonds (>75%). Species: *Tg – Toxoplasma gondii*; *Nc – Neospora caninum; Bb – Besnoitia besnoiti; Sn – Sarcocystis neurona; Ea/Ep/Et – Eimeria spp.; Cs – Cystoisospora suis; Cc – Cyclospora cayetanensis*.

### WNG kinases are secreted into the parasitophorous vacuole

As noted above, BPK1 has previously been identified as a PV-resident pseudokinase (30). We thus sought to assess the localization of other WNG kinases, and concentrated on the most divergent members of the family in *Toxoplasma*: ROP34 and ROP35 (Figure 1). We engineered parasite strains in which the endogenous copies of each of ROP34 and ROP35 were expressed in frame with a 3xHA tag. While both proteins appeared to be secreted into the PV, neither ROP35 nor ROP34 co-localized with the rhoptry marker ROP2 (Figure 2A). ROP35 co-localized well with the dense granular marker, GRA6, both within the parasites and after secretion into the PV (Figure 2B). ROP34 displayed a slightly different localization, in which the secreted protein localized to the basal end of the parasites within a vacuole. In addition, we observed brighter foci of ROP34 within parasites than within the vacuole, suggesting that ROP34 does not accumulate within the PV to the same extent as ROP35. While these data appear inconsistent with reported localization of ROP35 to the parasite PV via rhoptry secretion (32), we note that the previous report did not colocalize ROP35 with a known rhoptry marker, nor did it analyze endogenously tagged protein, both of which could lead to misinterpretation of the protein’s endogenous localization. As the “ROP” designation was originally created to indicate localization rather than function (33), we propose that the WNG kinases be renamed to avoid confusion with the unrelated ROPK family. Given its high conservation (Figure 1 and Supplemental Table S1c), we propose ROP35 be renamed WNG1, and other family members annotated as in Figure 1 and Supplemental Table S1c.

**Figure 2:**
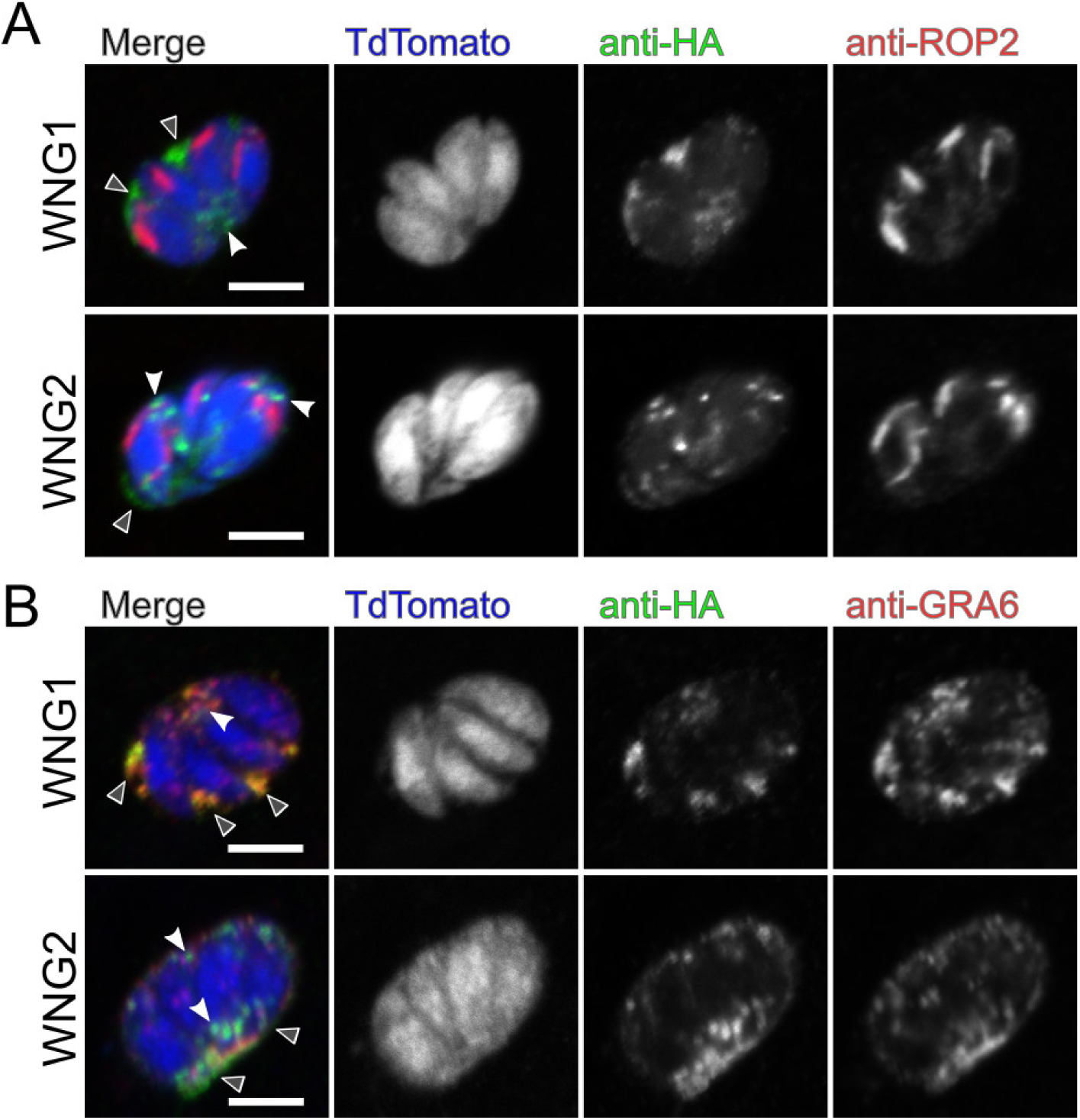
WNG kinases are secreted into the PV lumen from the dense granules. 0.5 μm confocal slices of WNG1-3xHA or WNG2-3xHA infected cells transiently transfected with TdTomato (blue) and stained with anti-HA (green) and either (A) the rhoptry marker anti-ROP2 (red) or (B) the dense granular/IVN marker anti-GRA6 (red). White arrowheads indicate intracellular signal; Gray arrowheads indicate secreted, PV-localized signal. Scale bars: 5 μm.

### The crystal structure of TgBPK1 reveals a non-canonical active site that lacks the Gly-loop

While the Gly-loop is thought to be both a critical catalytic and structural element of the protein kinase fold, a number of unusual kinases have been demonstrated to have either adapted a canonical kinase fold to perform a specialized non-catalytic function (34, 35, 11), or to use an atypical fold and active site to catalyze phosphoryl transfer (36, 37). We therefore sought structural information to better understand the topology of the WNG kinase fold. While we were unable to crystallize an active WNG kinase, we readily obtained crystals of the *Toxoplasma* pseudokinase BPK1 (Bradyzoite Pseudokinase 1). We solved the structure of BPK1 to 2.5 Å resolution (Figure 3A and Table 1). Like WNG1, BPK1 is secreted into the lumen of the PV (30), and is a clear member of the WNG family (Figure 1). As such, BPK1 shares both primary identity and predicted secondary structure with other WNG kinases throughout its sequence (Figure S1a), indicating that its structure would provide faithful insight into the WNG kinase fold.

**Table 1:** Crystallographic Data and Refinement.

|  | TgBPK1 (native) | TgBPK1 (Pt) |  |  |
| --- | --- | --- | --- | --- |
| **Data collection** |  |  |  |  |
| Space group | P2<sub>1</sub>2<sub>1</sub>2 | P2<sub>1</sub>2<sub>1</sub>2 |  |  |
| Cell dimensions |  |  |  |  |
| *a*, *b*, *c* (Å) | 171.47, 123.07, 86.62 | 184.77, 120.98, 92.36 |  |  |
| $\alpha$, $\beta$, $\gamma$ (°) | 90, 90, 90 | 90, 90, 90 | | |
|  |  | Peak | Inflection | Remote |
| Wavelength (Å) | 1.038 | 1.07195 | 1.07229 | 1.076276 |
| Resolution (Å) | 47.7 – 2.50 (2.59 – 2.50) | 50.0 – 3.75<br>(3.81 – 3.75) | 50.0 – 3.73<br>(3.79 – 3.73) | 50.0 – 3.92<br>(3.92 – 3.92) |
| Total reflections | 512518 | 130864 | 110497 | 105514 |
| R<sub>merge</sub> | 9.1 (86.3) | 8.3 (1.0) | 7.4 (88.2) | 8.6 (96.6) |
| CC<sub>1/2</sub> (final shell) | 0.76 | 0.63 | 0.70 | 0.75 |
| *I* / *I*σ | 23.8 (2.0) | 25.2 (1.8) | 25.3 (2.0) | 24.4 (2.2) |
| <sup>†</sup>Completeness (%) | 90.9 (63.5) | 90.3 (61.1) | 87.2 (63) | 89.8 (60.2) |
| Redundancy | 8.0 (7.3) | 6.0 (6.1) | 5.0 (5.1) | 6.0 (5.9) |
| **Refinement** |  |  |  |  |
| No. reflections | 58472 (4024) |  |  |  |
| R<sub>free</sub> reflections | 2987 (203) |  |  |  |
| R<sub>work</sub> / R<sub>free</sub> | 0.197 / 0.238<br>(0.238 / 0.289) |  |  |  |
| No. atoms |  |  |  |  |
| Protein | 12422 |  |  |  |
| Ligand/ion | 64 (EDO), 3 (CL) |  |  |  |
| Water | 381 |  |  |  |
| *B*-factors |  |  |  |  |
| Protein | 45.58 |  |  |  |
| Ligand/ion | 47.55 |  |  |  |
| Water | 33.12 |  |  |  |
| R.m.s. deviations |  |  |  |  |
| Bond lengths (Å) | 0.010 |  |  |  |
| Bond angles (°) | 1.37 |  |  |  |
| Ramachandran<br>(favored/disallowed) | 98.48 / 0 |  |  |  |
| Molprobity score | 1.14 |  |  |  |
| Molprobity clash<br>score | 3.48 |  |  |  |
| No. TLS groups | 20 per chain |  |  |  |

<sup>†</sup>Diffraction was anisotropic, which reduced the completeness, especially in highest resolution shell; for instance, for the native dataset diffraction was measured to 2.2 Å in the strongest dimension and 2.8 Å in the weakest.

**Figure 3:**
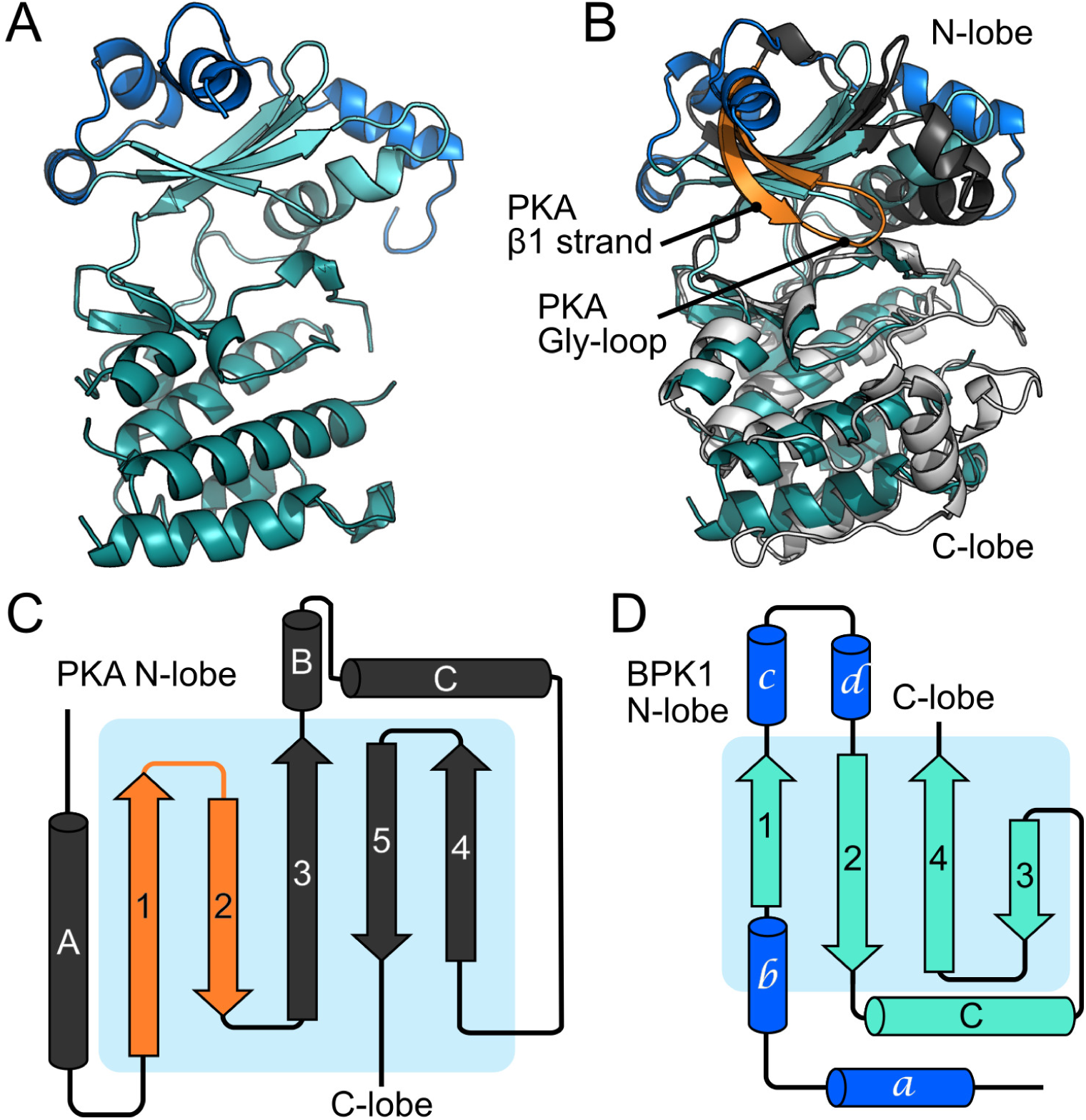
The structure of TgBPK1 reveals an atypical kinase fold lacking the Gly-loop. (A) Stereo view of the TgBPK1 structure. The N-lobe is in cyan, C-lobe colored in teal, and the helical “lid” that is unique to the WNG family is colored blue. (B) Superposition of the TgBPK1 structure with that of PKA (1ATP). TgBPK1 is colored as in (A). The N-lobe of PKA is dark gray, C-lobe is light gray, and β-strands that sandwich the Gly-loop are orange. Cartoon highlighting the differences between the N-lobes of (C) PKA and (D) TgBPK1, colored as in (B). Note the difference in the order of the N-lobe β-strands in PKA versus TgBPK1.

The BPK1 structure revealed a divergent kinase fold in which the Gly-loop and the first β-strand that stabilizes it (β1 in PKA nomenclature), have been replaced by a helical extension that packs against the top of the N-lobe of the kinase (Figure 3). Remarkably, not only do the WNG kinases lack a Gly-rich primary sequence, but have replaced the structural elements that compose the motif, resulting in a reorganized N-lobe architecture (Figure 3C,D). The core of the kinase fold, however, is remarkably well conserved, supporting our phylogenetic data (Figure 1) that suggest the WNG family diverged from a canonical Ser/Thr kinase fold. Two salt bridges help stabilize the BPK1 N-lobe within the pseudoactive site, including the bridge between the conserved αC-helix Glu and VAIK-Lys (Figure S3A). Notably, the lack of the Gly-loop and β1-strand creates an active site that is much more open than that of a canonical kinase, such as PKA (Figure S3B,C).

While BPK1 is a confirmed pseudokinase that cannot bind nucleotide (38), the other WNG kinase family members have conserved the other canonical motifs essential for catalysis (Figure S1a and Figure 4A, suggesting they may be active. To better understand how the WNG kinases have adapted their active sites to bind nucleotide and catalyze phosphoryl transfer without a Gly-loop, we modeled the WNG1/ROP35 active site using the structure of BPK1 as a template (Figure 4B,C). This model, together with analysis of sequence conservation among WNG1/ROP35 orthologs (Supplemental Figure S4), confirmed that the core of the canonical active site appears largely conserved (Figure 4C).

**Figure 4:**
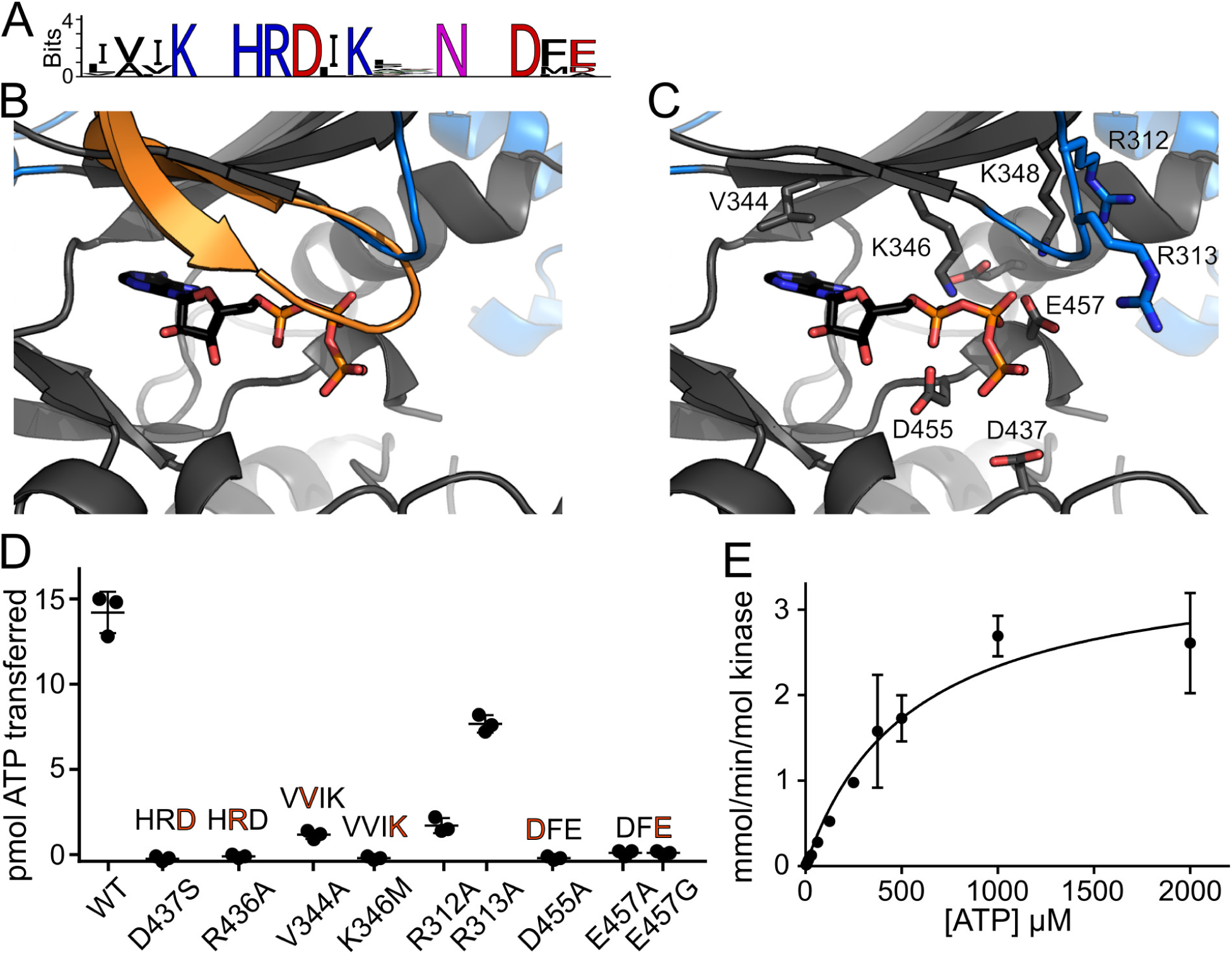
WNG1 has adapted its active site to catalyze phosphoryl transfer without a Gly-loop. (A) Sequence logos of the WNG kinase VAIK, HRD, and DFG motifs indicate conservation of critical catalytic residues. (B) A homology model of the WNG1 structure based on the BPK1 crystal structure, (gray and blue) has been superimposed with the structure of PKA (1.96Å backbone rmsd; 529 atoms compared). For clarity, only the PKA Gly-loop (orange) and bound nucleotide are shown. (C) A model of the WNG1 active site structure, colored as in (B). Bound ATP has been modeled based on superposition of the PKA structure. Residues that comprise either canonical motifs or WNG-specific substitutions are annotated and shown as sticks. (D) Kinase activities of wild-type WNG1 and the indicated mutant proteins using MBP as a protein substrate, quantified by ^32^P scintillation. Motifs altered by the mutants are shown above the data points. (E) A representative Michaelis-Menten fit of *in vitro* kinase assays of WNG1 using MBP as a substrate while varying ATP concentration.

We expressed and purified the kinase domain of *Toxoplasma* WNG1/ROP35 and found that it robustly phosphorylated the generic substrate MBP in an *in vitro* kinase assay. We verified that mutation of each of the canonical motifs that enable catalysis and Mg^2+^/ATP-binding (HRD*, VAIK*, D*FG) resulted in loss of kinase activity (Figure 4D). We also identified three notable variations from typical motifs within the active site. First, we noted that while substitution of the Ala in the VAIK motif to a bulkier side chain usually interferes with ATP-binding, WNG family members appear to prefer a Val at this position. Mutation of V344A in WNG1/ROP35 reduced the specific activity of the kinase to ~20% of wild-type (Figure 4D), consistent with a requirement for repositioning the ATP within the WNG active site. Second, we noted that WNG1/ROP35 orthologs have conserved a stretch of basic residues (R312/313 in *Toxoplasma*) that are placed near where the Gly-loop would lie (Figures 4B,C and S4). We therefore reasoned that the side chains of these residues may form a degenerate Walker A motif-like cap (28), and help replace the Gly-loop function. Consistent with such a model, both R312A and R313A mutants exhibited reduced specific activity, though R313A showed a much less severe effect than R312A (Figure 4D).

Finally, we noted that the WNG kinase Mg^2+^-coordinating DFG motif had an acidic residue (E447 in WNG1) replacing the Gly. As in our BPK1 structure (Figure S3A), the WNG1 E447 appears to form a salt-bridge with a conserved basic residue +2 from the VAIK Lys (Figure 4C; K348 in WNG1). This substitution is unusual for two reasons. (i) the DFG Gly is thought to be important for the regulation of many kinases, as it enables the peptide backbone to “flip” between two states (“DFG-in” and “DFG-out”; (39, 40)); (ii) the side chain of the Glu would be predicted to point towards the phosphates of the bound nucleotide (Figure 4C), and would thus electrostatically clash. We reasoned that a clash may be prevented, however, if the residue was participating in Mg^2+^-coordination, as the Asp in the DFG does. The pseudokinase domain of metazoan RNaseL also has this unusual substitution, in this case, a DFD motif. The crystal structure of RNaseL pseudokinase demonstrated that both acidic residues in the DFD motif participate in Mg^2+^-coordination (41), helping to explain the protein’s unusually high affinity (1 μM) for ATP. We therefore tested whether mutation of WNG1/ROP35 E457 to either Gly or Ala would affect its activity, and found that both mutant proteins had severely attenuated activity that was not significantly different from the kinase-dead HRD D437S mutant (Figure 4D).

We went on to determine that our recombinantly expressed WNG1 has an *in vitro* K_M,ATP_ of 520±90 μM (Figure 4E), using MBP as a substrate. Given the lack of the Gly-loop, which is a key ATP-binding element, it is unsurprising that this K_M,ATP,_ is higher than the 10-100 μM reported for many canonical kinases (42). However, the mammalian kinases Src and Akt have reported K_M,ATP_ of approximately 200 μM and 500 μM, respectively (42), indicating that our value for WNG1/ROP35 is consistent with an active kinase. Furthermore, the PV membrane is permeable to small molecules such as nucleotides (13, 14), and cellular ATP concentrations range between 2-5 mM (43), suggesting that PV nucleotide concentrations are well above that needed for activity with such an affinity for nucleotide.

Taken together, our structural and biochemical data suggest that WNG1/ROP35 and other family members are active protein kinases that have evolved multiple alterations to the active site to compensate for the lack of a Gly-loop. Furthermore, these broad structural changes imply an evolutionary pressure to reshape the protein structure to perform a specialized function.

### The intravacuolar network of parasites deficient in WNG1 kinase activity is unstable

We next sought to identify potential functions for the WNG kinases. We chose to concentrate our efforts on WNG1 because it is conserved throughout coccidia (Figure 1), concentrates within the PV lumen (Figure 2), and is important for chronic infection in a mouse model of infection (44). We used double homologous recombination to knock out the WNG1 locus in the RH*Δku80Δhxgprt* background (Supplemental Figure S5a). The resulting RH*Δwng1* parasites showed no obvious growth phenotype in normal culture conditions (not shown). We also generated WNG1-complemented strains by knocking a wild-type or kinase-dead (D437S; the HRD motif) copy of WNG1 into the empty Ku80 locus of the RH*Δwng1* strain. The kinase was expressed with its native promoter and in-frame with a C-terminal 3xHA. Both the active and kinase-dead complement strains expressed WNG1 at similar levels to the levels in the endogenously tagged parasite strain, and were appropriately localized to the dense granules and IVN (Supplemental Figure S5b).

To examine the ultrastructure within the vacuoles of parasites with and without active WNG1, we used transmission electron microscopy (TEM). We compared the vacuoles of HFFs that had been infected for 24 hours with either parental, RH*Δwng1*, or the complemented strains (Figures 5, S5c). The IVN is a complex structure of branching membranous tubules that fills a large portion of the PV lumen (45). As expected, we observed a dense network of tubules filling the lumenal space between the parental parasites (Figure 5A). While we did observe regions with IVN tubules in RH*Δwng1* vacuoles, they been largely replaced with unusual multilamellar structures containing many 70 – 150 nm diameter vesicles within a larger 0.5 – 2 μm membrane-delineated object (Figure 5B). These multilamellar structures appear much less electron dense than the tubular network, suggesting a lower protein content. Consistent with this observation, the internal vesicles appear to have been lost in some structures (Figure 5B, S5c), potentially due to reduced crosslinking before plastic embedding. Importantly, we prepared samples from mutant parasite strains in parallel with a parental control. We never observed loss of tubular structures in the parental strains, suggesting that this phenotype is not an artifact of our preparation. While vacuoles of wild-type WNG1 complemented parasites were indistinguishable from the parental, those formed by the kinase-dead complemented strain exhibited the same loss of IVN tubules and its apparent replacement with large multilamellar vesicles (Figures 5C-D). These changes were quantified in Figure 5E,F. These data indicate that WNG1 phosphorylates one or more proteins involved in IVN biogenesis and stability, and that this phosphorylation is required for normal function.

**Figure 5:**
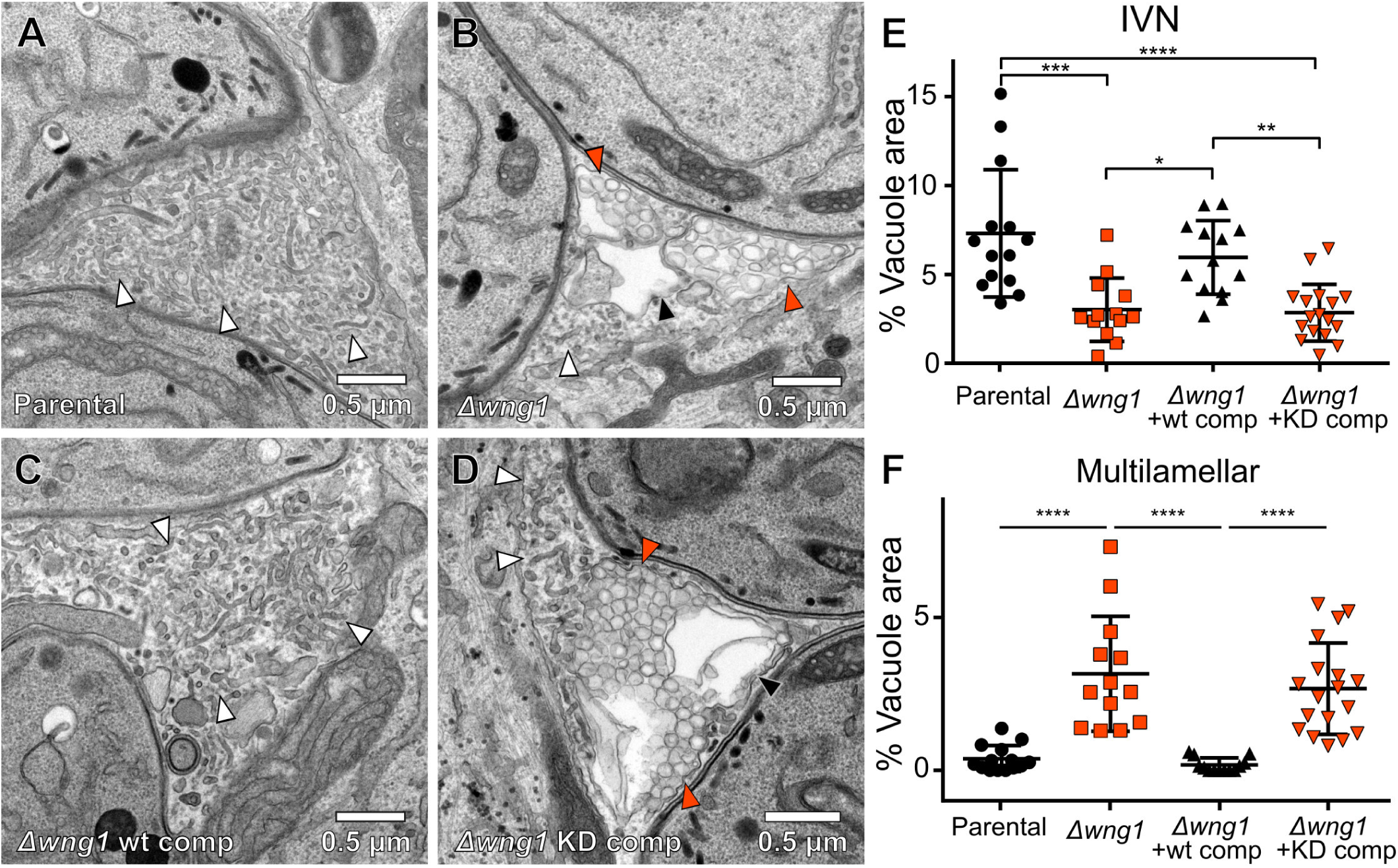
Vacuoles lacking active WNG1 kinase show disrupted IVN membranes. Representative transmission electron microscopic images of the (A) Parental, (B) RH*Δwng1*, (C) RH*Δwng1* complemented with wild-type WNG1, and (D) kinase-dead complemented strains. IVN tubules are indicated with white arrowheads. Multilamellar vesicles are indicated with solid orange arrowheads. Multilamellar structures in which internal vesicles appear to have been lost during fixation are indicated with a black arrowhead in (B) and (D). The relative area of each IVN tubules and multilamellar vacuole from EM images as in (A-D) were quantified in ImageJ. Significance was calculated in Prism by ANOVA; p<0.0001 (****); p<0.001 (***); p<0.01 (**); p<0.05(*).

### Quantitative phosphoproteomics reveals GRA proteins as candidate substrates of WNG1

To identify potential substrates of WNG1, we compared phosphoproteomes of the parental (WT) and RH*Δwng1* strains using stable isotope labeling with amino acids in cell culture (SILAC) quantitative mass spectrometry (MS) based proteomics as previously described (46). Briefly, we infected human foreskin fibroblasts (HFFs) for 24 h with WT or RH*Δwng1* parasites previously grown in either “heavy” (H) or “light” (L) SILAC media. After cell lysis we mixed the samples (H and L) in 1:1 ratio applying forward (*Δwng1*/WT), reverse (WT/*Δwng1*) as well as control labeling (WT/WT). This mixing strategy ensures that both systematic and technical errors due to stable isotope labeling can be identified and results in high confidence of MS quantifications. Mixed lysates were then digested with LysC/trypsin and phosphopeptides enriched and fractionated as described in the methods section. We prepared 3 biological replicates for WT vs Δwng1 samples and analyzed quantitative differences in the proteome and phosphoproteome between WT and mutant samples by mass spectrometry. We identified 10,301 phosphosites for both human and *Toxoplasma* and obtained quantification (H/L ratios) for 8,755 of them. *Toxoplasma*-specific sites constituted 2,296 (~30%) of all quantified sites (Supplemental Table S6), which is a similar proportion of sites identified in previous studies using intracellular *Toxoplasma* parasites (23). In order to identify significantly changing sites between WT and Δwng1 parasites a one sample t-test was performed applying the following parameters: p-value < 0.05 and |log_2_| fold change > 1 (Figure 6). Furthermore, phosphosite significance was also correlated with the SILAC control sample (WT/WT) and the proteome data to control for differential phosphorylation originating from the technical variation in the system and protein abundance, respectively (Supplemental Table S6). We also identified a number of phosphorylation sites on protein with consistent loss of phosphorylation in RH*Δwng1* parasite strains, that however, did not pass the t-test significance test (Supplemental Table S6). However, all phosphorylation sites close to the p-value cutoff are predicted or known secreted proteins, indicating that the p-value may be overly stringent in this case.

**Figure 6:**
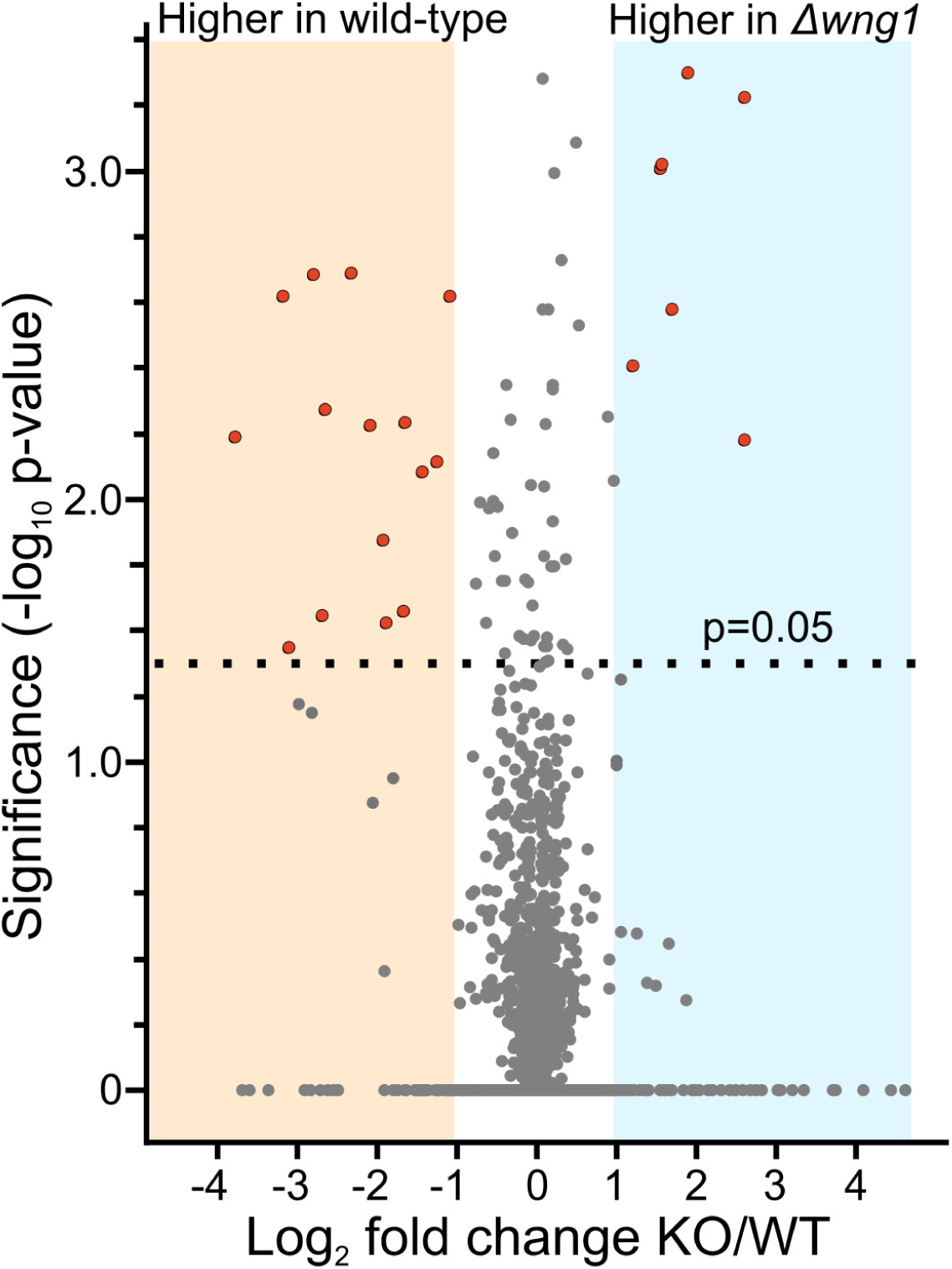
Overview of quantitative phosphoproteomics data. SILAC quantification of change in phosphosite abundance plotted against significance of change for 2296 phosphosites in RH*Δwng1* versus parental parasites. See Supplementary Table 5 for full data set. Significantly changing phosphosites (p-value < 0.05 and −1 > log_2_ change > 1) enriched in dense granule proteins are highlighted.

We identified 10 proteins in which phosphorylation was significantly reduced between the parental and RH*Δwng1* samples (Figure 6, Table 2, and Supplemental Table S6). Among these candidate substrates were 6 proteins well-known to be associated with the parasite IVN tubules (Table 2), including GRA2 and GRA6, which are essential for IVN biogenesis (45). Another hit, GRA37, was identified in a recent proteomics analysis of PV membrane proteins, and was found to colocalize with IVN markers (47). We identified three proteins with WNG1-dependent phosphorylation that have not been previously studied, and are therefore annotated as “hypothetical” in the genomic database (ToxoDB v32 gene models: TGGT1_244530, TGGT1_254000, and TGGT1_267740). We reasoned that if WNG1 is, indeed, a PV-resident kinase, WNG1-dependent phosphorylation should predict PV (and possibly IVN) localization. We therefore engineered strains in which the proteins were endogenously tagged at their C-terminus with a 3xHA epitope. Immunofluorescence revealed that each of these proteins were secreted into the PV, and co-localized with the dense granular and IVN marker GRA6 both within parasites and within the vacuolar lumen (Figure 7A). We have thus annotated these three genes as encoding newly described dense granular proteins GRA43, GRA44, GRA45 (Tables 2 and 3).

**Table 2:**
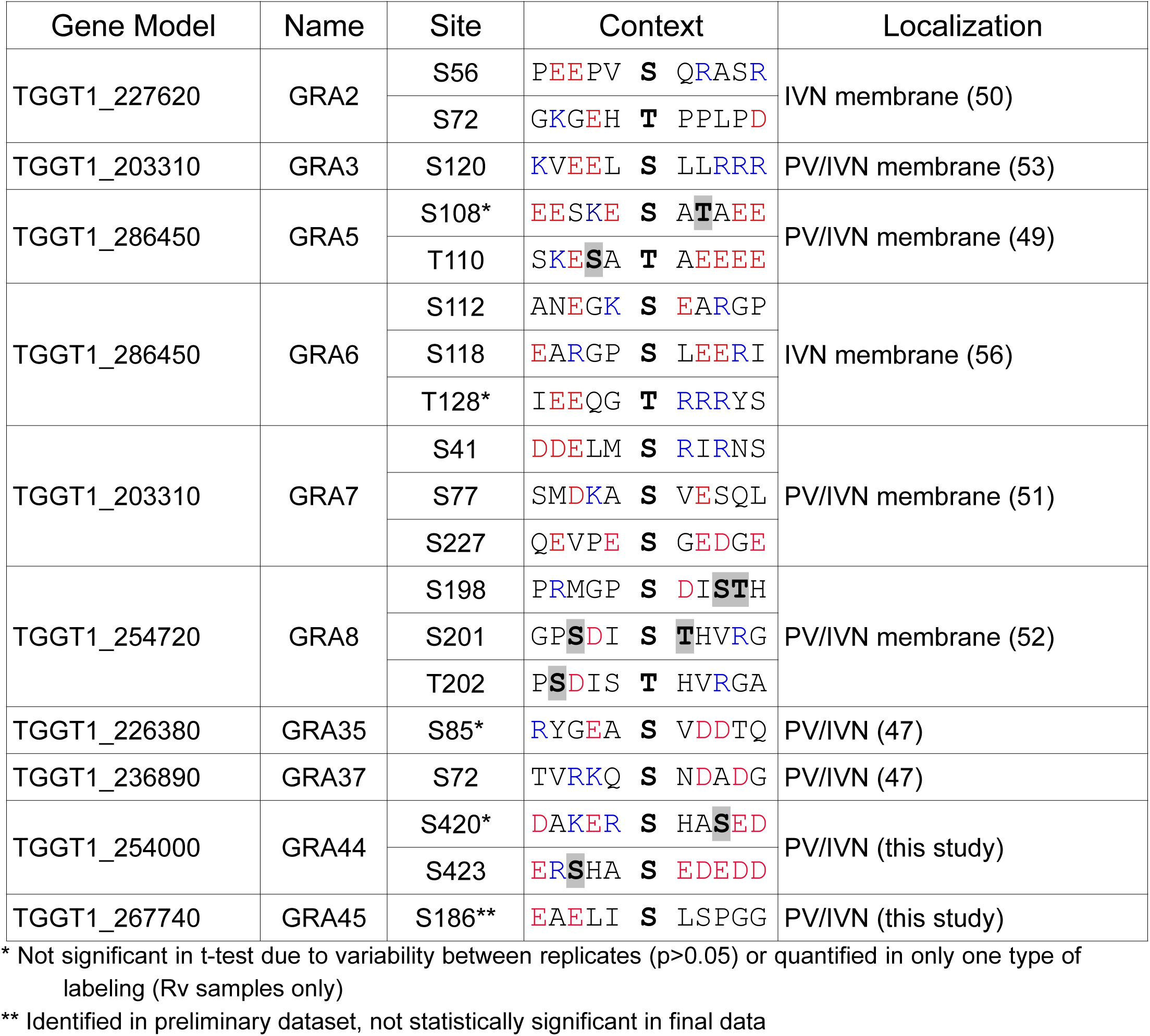
Phosphosites lost in RH*Δwng1* vacuoles. The sequence context of each of the phosphosites is indicated. Acidic residues are red, basic residues are blue. Note that some regions appear to be hyperphosphorylated in a WNG1-dependent manner. Such potential priming sites are indicated bolded with a gray background in the phosphosite context.

**Table 3:**
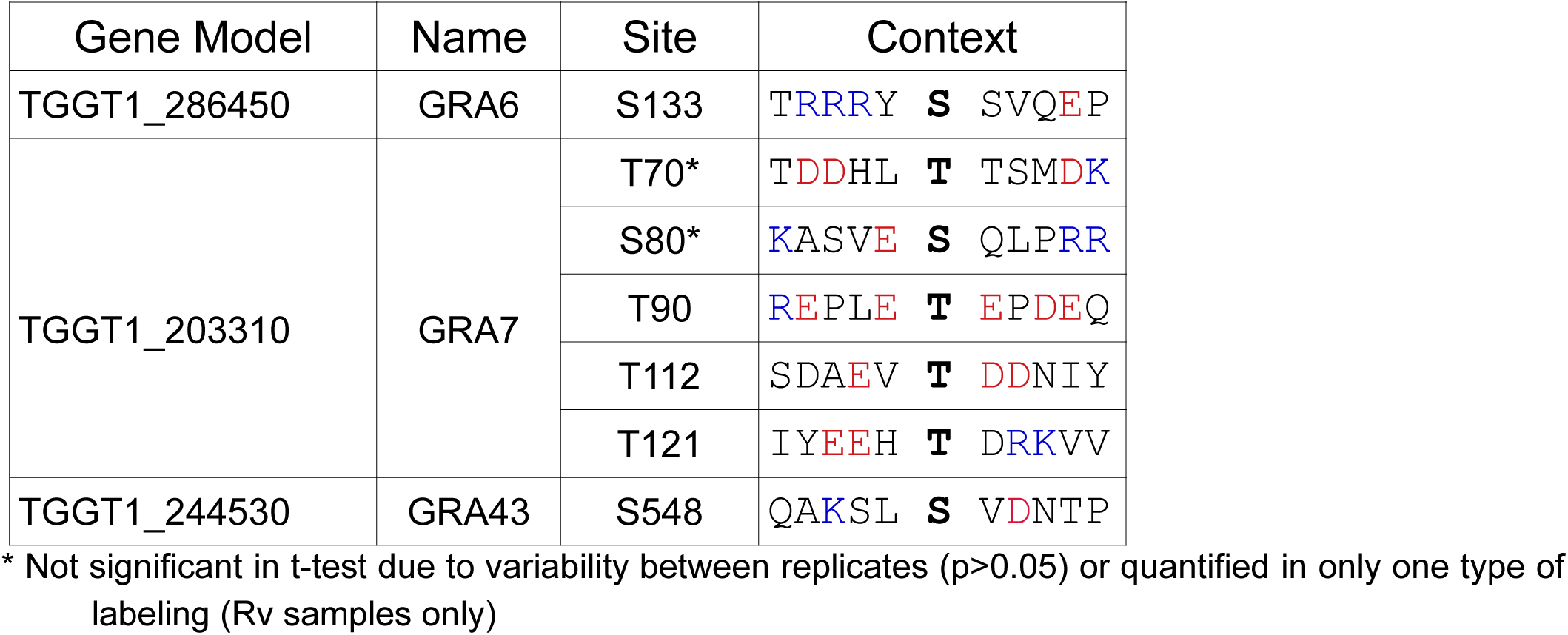
Phosphosites upregulated in WNG1-deficient vacuoles. Table is formatted as in Table 2.

**Figure 7:**
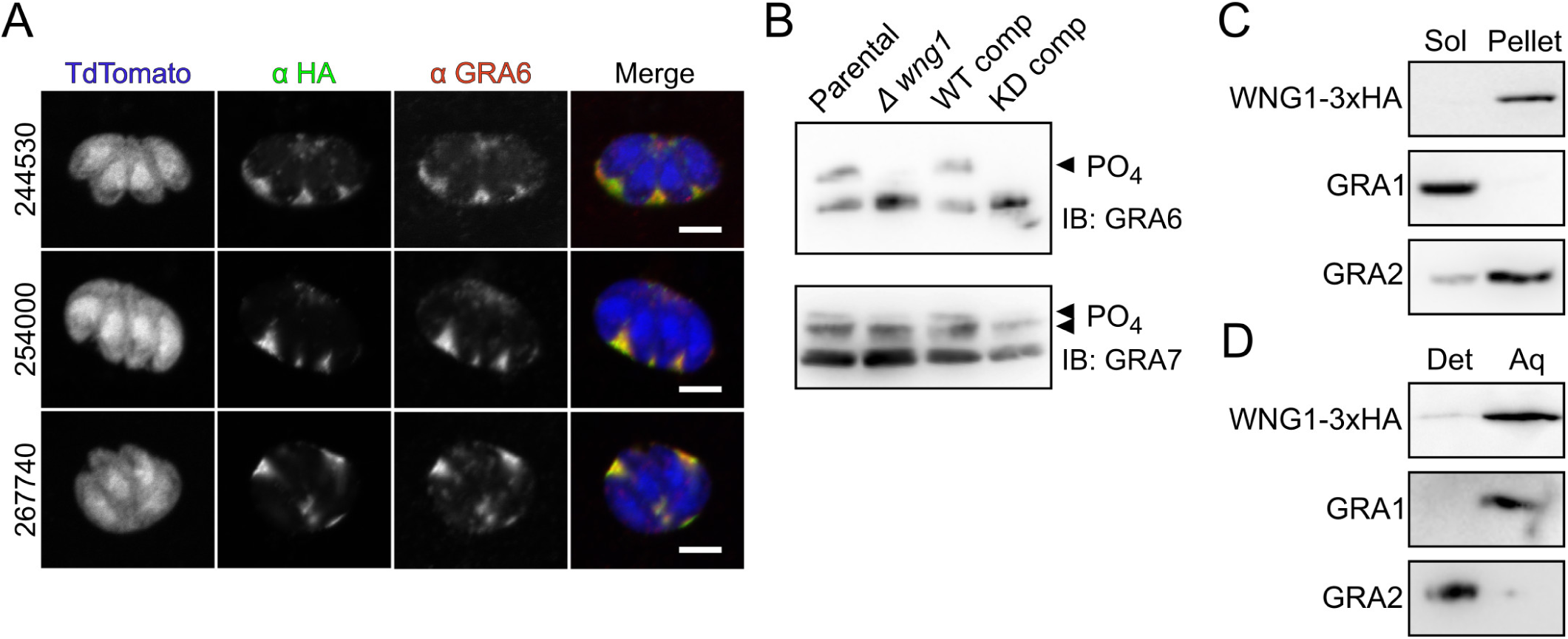
WNG1 and its substrates are membrane associated. (A) 0.5 μm confocal slices of cells infected with parasites in which the indicated unannotated candidate substrates are endogenously 3xHA tagged (anti-HA; green), transiently transfected with TdTomato (blue), and co-stained with the dense granular and IVN marker GRA6 (red). Scale bars 5 μm. (B) Western blot of lysates of cells infected with the indicated wild-type, knockout, or complement strains probed with anti-GRA6 and anti-GRA7 antisera. Phosphorylated bands are indicated with arrowheads. (C) Western blot of host and PV membranes that have been ultracentrifuged. WNG1-3xHA is detected with anti-HA, GRA1 and GRA2 are used as soluble and membrane-associated controls, respectively. (D) Western blot of host and PV membranes that have been subjected to Triton-X-114 partitioning between detergent (Det) and aqueous (Aq) phases.

In addition to the phosphosites that were lost in the vacuoles lacking WNG1, we identified 7 sites where phosphorylation was significantly increased in the RH*Δwng1* samples over parental (Table 3). These include 1 site on GRA6, 5 sites on GRA7, and 1 site on TGGT1_244530. The phosphorylated states of GRA6 and GRA7 in cells infected with wild-type parasites are readily distinguishable by SDS-PAGE and western blot (16, 48). To confirm the changes in phosphorylation of these proteins, we blotted lysates of cells infected with either the parental or RH*Δwng1* strains (Figure 7B). To demonstrate that these changes were due to the presence of WNG1, and to confirm the requirement of WNG1 kinase activity, we also assessed GRA protein membrane association in the wild-type and kinase dead WNG1 complemented strains (Figure S5a). We then analyzed lysates of cells infected with each the above strains by western blot probed with either anti-GRA6 or anti-GRA7 antibody. The slower migrating, phosphorylated band of GRA6 was apparent in both parental and wild-type complemented lysates, but was undetectable in the knockout and kinase-dead complemented lysates (Figure 7B). Consistent with complex WNG1-dependent differences in phosphorylation of GRA7, we observed a reduction in the total amount of phosphorylated GRA7 in the knockout and wild-type complemented parasites, but observed other slowly migrating bands, presumably the novel phospho-states listed in Table 3. Our data thus demonstrate WNG1 is an active, PV-resident kinase that is required for the phosphorylation of proteins associated with the PV-facing leaflet of the IVN membrane.

Given that these candidate WNG1 substrates have been demonstrated to either associate with or integrate into PV membranes (48–53), we asked whether WNG1 itself was membrane associated once secreted into the PV. To test this, we mechanically disrupted a human foreskin fibroblast (HFF) monolayer that had been highly infected with WNG1-3xHA parasites. Intact parasites were separated from host and PV membranes by a low speed (2400 g) spin, and the resulting supernatant was further separated by ultracentrifugation. WNG1, like the known integral membrane protein (and putative WNG1 substrate) GRA2, was found largely in the membrane-associated pellet (Figure 7C). In parallel, we partitioned an aliquot of the same low speed supernatant with Triton-X-114 (54). While GRA2 was found entirely in the detergent phase, WNG1 partitioned in the aqueous phase (Figure 7D), indicating that it is a soluble protein that is membrane-associated, rather than integrating into the membrane directly. Such a non-integral association of WNG1 with the PV membrane is consistent both with the lack of a predicted transmembrane, amphipathic helix, or other membrane association domain in the WNG1 sequence, and with our ability to purify soluble recombinant protein.

### Efficient membrane association of proteins involved in intravacuolar network biogenesis depends on WNG1 kinase activity

The trafficking of IVN-associated proteins is highly unusual. Many GRA proteins integrate amphipathic or transmembrane helices into the IVN membrane, but remain soluble while trafficking through the parasite secretory system (48, 49, 53), presumably by complexing with an unidentified chaperone. Notably, many of the WNG1-dependent phosphorylation sites are located in predicted helical regions of sequence (Figure S8A) that have been shown to be required for GRA membrane association (50, 55). We therefore reasoned that phosphorylation of substrates by WNG1 may help regulate the switch from soluble to membranous states of IVN-associated GRA proteins. To test this hypothesis, we assessed WNG1 membrane association by comparing fractionated lysates from parental, RH*Δwng1*, and the kinase-active and kinase-dead complement strains. We prepared samples from 6 independent infections per condition, which were then separated by SDS-PAGE and analyzed by protein immunoblotting using antibodies recognizing various GRA proteins as indicated in Figure 8. We observed no difference in Triton-X-114 partitioning for any of the strains (Figure S8B). We quantified the relative soluble amounts of each protein (Figure 8B), which revealed a requirement for WNG1 kinase activity on IVN GRA membrane association. In particular, GRA4, GRA6, and GRA7 exhibited significant reductions in the fraction of protein that was PV membrane-associated in the RH*Δwng1* and kinase-dead samples. Notably, the phosphorylated forms of GRA6 and GRA7 are found exclusively in the membrane-associated fractions (Figure 8A), and the loss of the phosphorylated states does not appear to result in a concomitant increase in unphosphorylated protein at the membrane. Taken together, our results suggest that WNG1-dependent phosphorylation of the GRA proteins promotes their association and is critical for the proper formation of the IVN.

**Figure 8:**
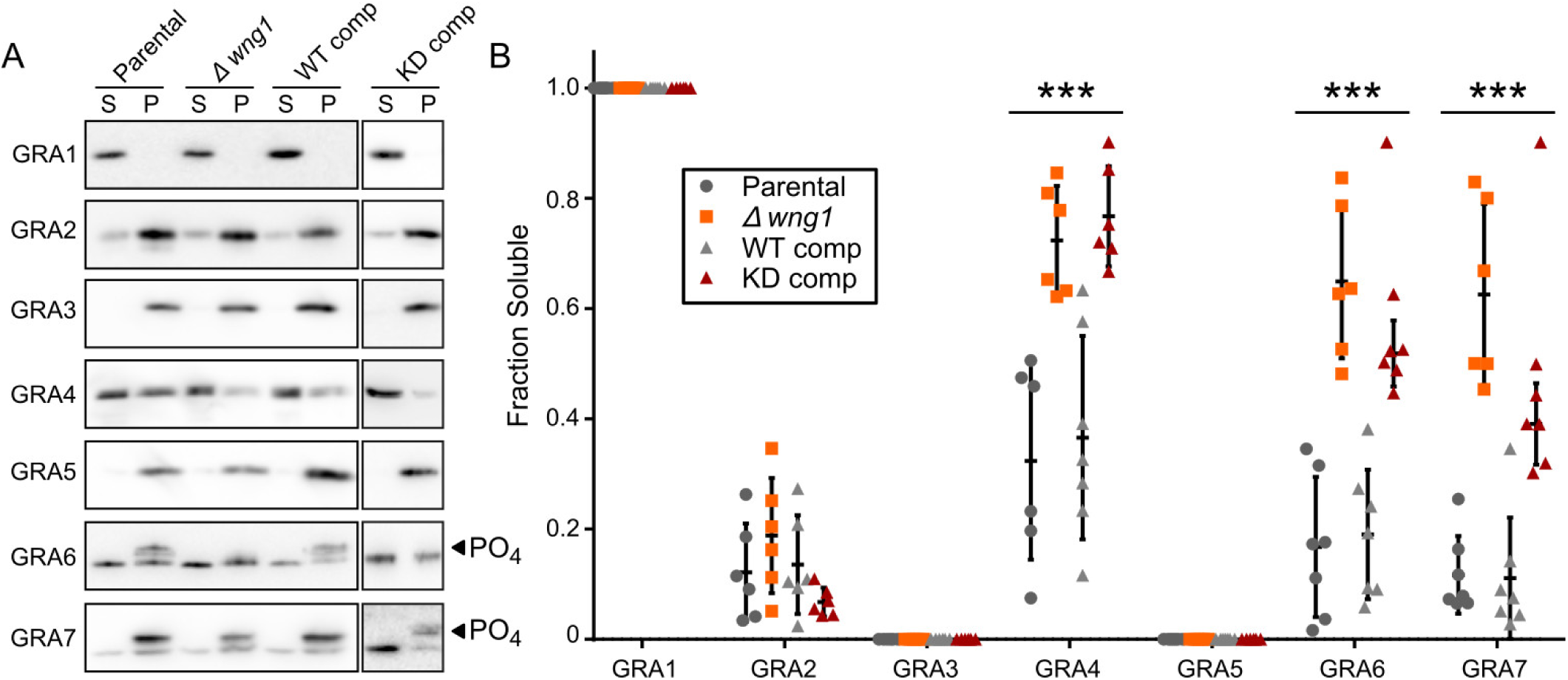
Membrane association of GRA proteins correlates with WNG1 kinase activity. (A) Representative western blot of samples in which host and PV membranes were ultracentrifuged and the soluble (S) and pellet (P) fractions were separated by SDS-PAGE and probed with the indicated antisera. Phosphorylated bands are indicated with arrowheads. (B) Quantification of n=6 biological replicates as in (A). Significance was calculated by ANOVA in Prism; p<0.001 (***).

## Discussion

We have identified an unusual family of parasite-specific protein kinases that divergently evolved from a canonical protein kinase fold and have lost the typical Gly-rich loop. We have demonstrated that, in spite of missing a structural element thought to be critical to nucleotide binding and catalytic activity, the WNG kinases can catalyze phosphoryl transfer. Through structural and biochemical analyses, we have delineated subtle changes to the kinase active site that facilitate its catalytic activity. We went on to show that the most conserved member of the family, WNG1/ROP35, is secreted by *Toxoplasma* into the PV, where it associated with the PV membranes. We found that WNG1 kinase activity is required for the phosphorylation of many of the proteins known to be associated with the IVN membranes. Furthermore, loss of WNG1 kinase activity was correlated with a reduction in membrane association for a subset of the IVN proteins for which there are antibodies available. Finally, we found that parasite vacuoles deficient in catalytically active WNG1 have a substantial reduction in their IVN, suggesting that kinase activity is required for either the efficient formation or stability of the IVN membrane tubules.

The unusual WNG kinase fold raises the question: what may have been the evolutionary pressure that drove the divergence of the WNG family and loss of the Gly-loop? WNG1 is the most conserved member of the family, and appears to preferentially phosphorylate sites on proteins closely associated with the IVN membrane. Moreover, many of the sites we identified are at or near predicted helices (Figure S8A) that have been previously implicated in GRA protein interaction with membranes (50, 55, 56), or, in the case of GRA3, within a predicted coiled-coil. The rearrangement of the WNG active site has resulted in an unusually open active site (Figure S3B,C) that may better accommodate such folded or otherwise sterically restricted substrates. The atypical “alpha” family of kinases (57) are also able to phosphorylate helical substrates, such as the coiled-coil domains of myosin heavy chains (58). The alpha kinases share no detectable sequence homology to canonical protein kinases in spite of their similar overall folds (59, 60). The active sites of alpha kinases differ in several ways from canonical protein kinases. As with the WNG kinases, the alpha kinases have a more open active site that would accommodate a helical substrate (60). In any event, a comprehensive understanding of the mechanisms of substrate recognition in atypical kinases such as the WNG and alpha kinase families will require structural studies of kinase:substrate complexes.

Notably, the phosphosites we identified as WNG1-dependent are not detectable in extracellular parasites (23), indicating that WNG1 phosphorylates its substrates in the PV lumen rather than while trafficking through the parasite secretory system. Our phosphoproteomics data revealed both phosphosites lost in WNG1 knockout parasites, as well as a smaller number of upregulated sites that were only detectable when WNG1 was missing. These data suggest that another kinase is capable of phosphorylating a subset of sites on the IVN GRA proteins, and its activity may be partially compensating for WNG1 loss. Alternatively, this other kinase activity may be acting in competition with that of WNG1. It is possible that these novel sites are phosphorylated by another member of the WNG family. Regardless, the data we present here are consistent with a role for WNG1-mediated phosphorylation in the regulation of protein-protein and/or protein-membrane interactions of PV-resident proteins.

The multilamellar vesicles we observe in vacuoles deficient in WNG1 kinase activity are reminiscent of structures that have been previously observed during the first steps of IVN biogenesis (15). While non-phosphorylated, recombinantly-expressed GRA2 and GRA6 are sufficient to tubulate large unilamellar vesicles *in vitro* (19), it is possible that WNG1 kinase activity is required to ensure the efficiency of this process in cells. This may be explained by an apparent paradox that exists in GRA protein trafficking: GRA proteins that are destined to integrate into PV membranes traffic through the parasite secretory system as soluble entities (48–50), presumably in complex with an unknown solubilizing protein (Figure 9). Such a switch ensures that the parasite’s intracellular and plasma membranes are protected from the tubulating activity of the GRA proteins. Removal of a solubilizing chaperone normally requires energy provided by ATP hydrolysis. There are no known chaperones secreted into the *Toxoplasma* PV. Consistent with a model in which WNG1 regulates membrane association of a subset of PV GRAs, we observed that each GRA4, GRA6, and GRA7 had were substantially more soluble in the vacuoles of parasites deficient in WNG1 kinase activity. There is thus an intriguing possibility that the ATP used during WNG-mediated phosphorylation is providing the energy to drive membrane insertion of a subset of GRA proteins (Figure 9). Such a non-canonical chaperoning mechanism is not without precedent. The mammalian neuropeptide 7B2 solubilizes the prohormone convertase 2 as it traffics to the Golgi (61), where 7B2 is phosphorylated by a resident kinase, resulting in release of the complex (62, 63).

**Figure 9:**
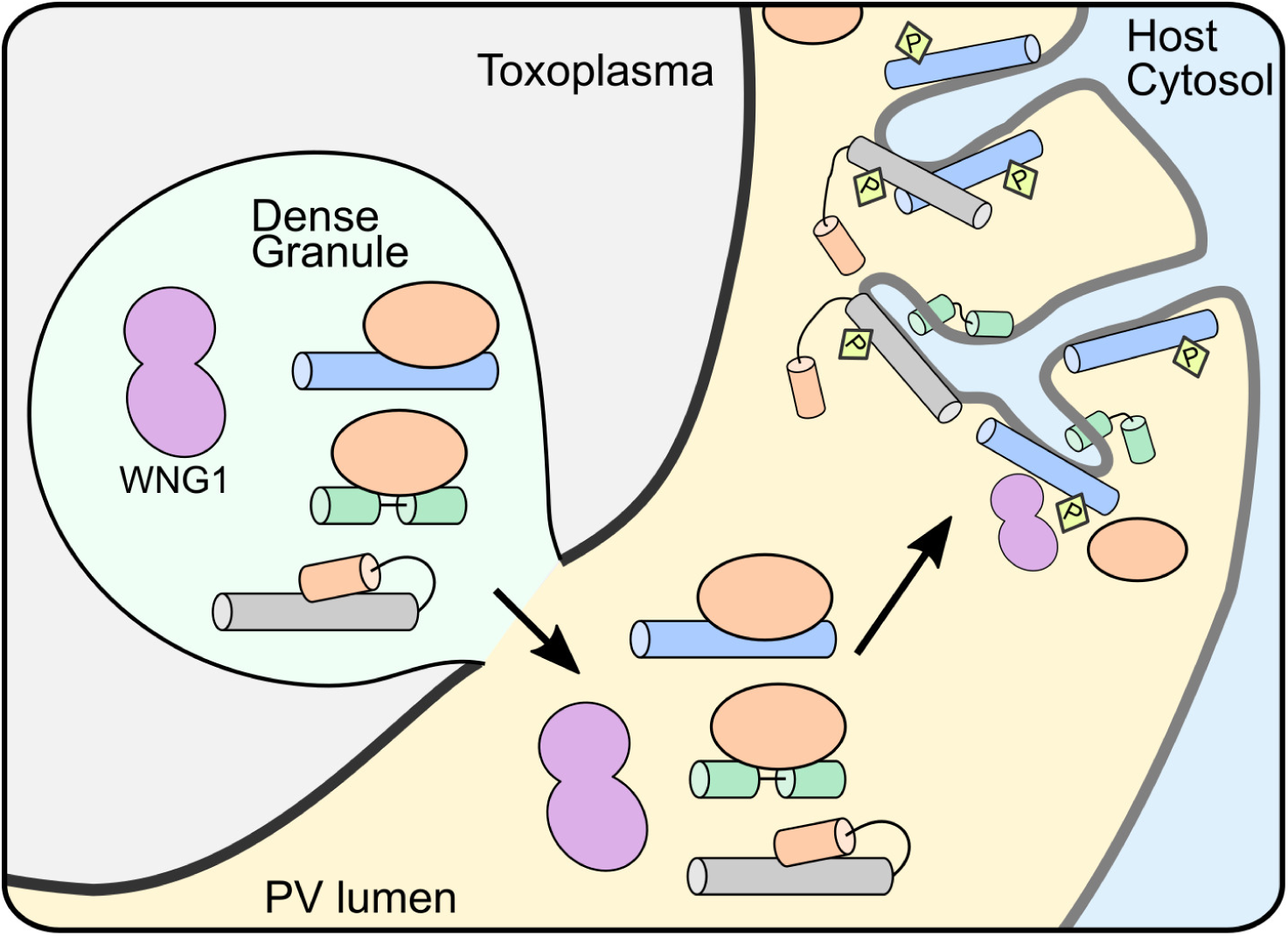
Model for WNG1 regulation of IVN GRA protein membrane association. Within the parasite secretory pathway, membrane-seeking GRA proteins (blue, green, and gray cylinders) are complexed with solubilizing proteins or domains (orange). Once secreted into the PV lumen, WNG1 is activated through an unknown mechanism and phosphorylates the GRAs, leading to their eventual insertion into the PV membrane and efficient stabilization of the IVN tubules.

In spite of decades of study, the *Toxoplasma* parasitophorous vacuole remains a mysterious organelle. The major function identified for IVN-associated proteins is in IVN biogenesis, as the deletion of either GRA2 or GRA6 results in a complete loss of the structure (45). Such IVN-deficient parasites have been used to link the IVN to nutrient uptake (16–18) and immune evasion (19, 21), though the precise mechanisms and roles of the IVN have not been established in these processes. Consistent with the pleiotropic effects of disrupting the IVN, knockout of IVN-associated proteins strongly attenuates parasite virulence (22, 64). Infection of mice with WNG1 knockout parasites yields a substantially reduced cyst burden (44), which is consistent with the role we observed for WNG1 in IVN biogenesis and stability and the likely resulting pleiotropic effects on the parasite’s biology. Our discovery of potential regulatory phosphorylation may facilitate future work to associate specific GRA protein complexes with their biochemical functions and thus better delineate the roles of the IVN in parasite pathogenesis.

## Materials and Methods

### Phylogenetic analysis

Protein sequences for the WNG kinases were identified by using custom scripts that iteratively BLAST (65) the ToxoDBv24 collections for *Toxoplasma gondii*, *Neospora caninum*, *Sarcocystis neurona*, *Eimeria spp.*, *Cystoisospora suis*, and *Cyclospora cayetanensis*. *Besnoitia besnoiti* sequences from the Uniprot nonredundant collection. Multiple sequence alignments were generated by MAFFTv7 (66), and manually edited as necessary. The maximum likelihood phylogenetic tree and bootstrap analysis (1000 replicates) were estimated using RAxML v8.1.17 (67), and the resulting tree was annotated using a script based on the jsPhyloSVG package (68) and Inkscape.

### PCR and plasmid generation

All PCR was conducted using Phusion polymerase (NEB) using primers listed in Supplemental Table S10. Constructs were assembled using Gibson master mix (NEB). Point mutations were created by the Phusion mutagenesis protocol.

### Parasite culture and transfection

Human foreskin fibroblasts (HFF) were grown in Dulbecco’s modified Eagle’s medium supplemented with 10% fetal bovine serum and 2 mM glutamine. *Toxoplasma* tachyzoites were maintained in confluent monolayers of HFF. Epitope-tagged and knockout parasites were generated by transfecting the RH*Δku80Δhxgprt* strain (69) with 15 μg of linearized plasmid and selecting for HXGPRT expression, as previously described (70). The loxP-flanked HXGPRT selection cassette in knockout parasites was removed by transient transfection with a plasmid overexpressing Cre recombinase, and selecting with 6-thioxanthine. WNG1 complement parasites were created by targeting 3xHA-tagged WNG1 (either wild-type or kinase-dead) driven by its native promoter, together with a bleomycin resistance cassette, to the empty Ku80 locus, and selecting with bleomycin, as previously described (71).

### Immunofluorescence

HFF cells were grown on coverslips in 24-well plates until confluent and were infected with parasites. The cells were rinsed twice with phosphate buffered saline (PBS), and were fixed with 4% paraformaldehyde (PFA)/4% sucrose in PBS at room temperature for 15 minutes. After two washes with PBS, cells were permeabilized with 0.01% Triton-X-100 for 10 minutes and washed 3x with PBS. After blocking in PBS + 3% BSA for 30 min, cells were incubated in primary antibody in blocking solution overnight at room temperature. Cells were then washed 3x with PBS and incubated with Alexa-fluor conjugated secondary antibodies (Molecular Probes) for 2 h. Cells were then washed 3x with PBS and then mounted with mounting medium containing DAPI (Vector Laboratories). Cells were imaged on a Nikon A1 Laser Scanning Confocal Microscope. Primary antibodies used in this study include rat anti-HA (Sigma; 1:500 dilution), mouse anti-GRA2 (BioVision; 1:1000 dilution), mouse anti-GRA6 (gift of David Sibley; 1:1000 dilution), rabbit anti-ROP2 (1:10,000 dilution).

### Protein purification

BPK1 (residues 61-377 cloned into pGEX4T) was expressed as a GST fusion in *E. coli* Rosetta2(DE3) overnight at 16°C after induction with 300 mM IPTG. Cells were resuspended in 50 mM Tris 8.0, 200 mM NaCl, 1% Triton-X-100 and 0.2% sodium sarkosyl, lysed by sonication, and centrifuged at 27k rcf for 30 min. GST-fusion protein was affinity purified using glutathione sepharose, which was washed first with PBS containing 1% Triton-X-100, and then without detergent. Protein was eluted by overnight on-bead thrombin cleavage at 4°C overnight. BPK1 was further purified by anion exchange and size exclusion chromatography, where it was flash frozen in 10 mM HEPES, pH 7.0, 100 mM NaCl for storage. Recombinant wild-type and mutant WNG1 (residues 265-591; cloned into pET28) proteins were expressed N-terminally fused with His_6_-SUMO in *E. coli* Rosetta2(DE3) incubated overnight at 16°C after induction with 300 mM IPTG. Bacteria were lysed in 50 mM HEPES 7.4, 500 mM NaCl, 15 mM Imidazole, lysed by sonication, and centrifuged as above. His_6_-fusion proteins were affinity purified using NiNTA resin, and eluted in 50 mM Tris, pH 7.0, 500 mM NaCl, 250 mM imidazole and dialyzed in 20 mM Tris, pH 7.0, 300 mM NaCl before concentration and flash freezing for long-term storage.

### Protein crystallization

Small hexagonal plates of BPK1 grew in a wide variety of conditions in initial screens. High quality crystals were seeded from initial hits grown in 0.2M Proline, 0.1M HEPES 7.4, 10% PEG-3350. To generate a platinum derivative, crystals were soaked with reservoir solution containing 10 mM K**_2_**PtCl_4_ for 2 h and washed quickly in reservoir solution. All crystals were flash frozen in a cryoprotectant of reservoir with 25% ethylene glycol.

### Data collection, structure determination, and refinement

The diffraction data for the native crystals were collected at beamline 19-ID at the Advanced Photon Source at a wavelength of 1.038 Å and a temperature of 100 K. Native crystals diffracted to 2.5 Å, though diffraction was highly anisotropic, ranging from 2.2 Å in the best dimension to 2.8 Å in the worst. Data for the platinum dervatives were collected in an inverse beam experiment at 1.07195 Å, 1.076276 Å, and 1.07229 Å, corresponding to peak, remote, and inflection wavelengths. Integration, indexing, and scaling of the diffraction data were performed using the HKL2000 suite of programs (72). Initial phases at 3.5 Å were determined by multiwavelength anomalous diffraction from the Pt datasets using the SHELX suite (73) and used to generate a starting model after density modification with the SOLVE/RESOLVE package (74, 75). The high resolution native data were incorporated for extension and map improvement in Phenix (76). Manual rebuilding in Coot (77) and refinement in Refmac5 (78), led to a final 2.5 Å structure of BPK1 (PDB accession: 6M7Z). The structure was evaluated with Molprobity (79).

### Homology modeling

A model of the WNG1/ROP35 structure was created in Modeller v9.14 (80) using the BPK1 structure as a template and an alignment of BPK1 and WNG1 created using Clustal Omega (81).

### In vitro kinase assays

The kinase assays comparing WT and mutant activities were run using 2 µM of His_6_sumo-WNG1, 4 mM MgCl2, 200 µM cold ATP, 1mM DTT, 1mg/mL BSA, 10% glycerol, 300 mM NaCl, 20 mM Hepes pH 7.5. Reactions were started by adding a hot ATP mix, that contained 10 µCi ϒ[^32^P] ATP and 5 µg MBP. The K_M,ATP_ kinase assays were run using the same mix as above except non-radioactive ATP was used in a range of concentrations from 1 mM to 32.5 µM. The 25 µL reactions were incubated at a 30°C water bath for 2 h. Reactions were stopped by adding 9 µL 4x SDS-buffer. 20 µL samples were then run on an SDS-PAGE gel. The gels were coomassie stained, the MBP band, excised and radioactivity quantified using a scintillation counter. All data were analyzed using GraphPad Prism 7.

### Cell culture, lysis and protein digestion for MS proteomics

All reagents were obtained from Sigma-Aldrich unless specified otherwise. Parental (WT) and RH*Δwng1 Toxoplasma* parasites were cultured in either R0K0 (light) or R10K8 (heavy) SILAC medium (Dundee Cell Products) for 8 generations to ensure efficient heavy label incorporation. 24 h prior cell lysis human foreskin fibroblasts (HFFs) were infected (MOI=5) with WT or RHΔwng1 parasites. Lysis was then performed in 8 M urea, 75 mM NaCl, 50mM Tris, pH=8.2, supplemented with protease (complete mini tablets, Roche) and phosphatase (Phos Stop tablets, Roche) inhibitors followed by sonication to reduce sample viscosity (30 % duty cycle, 3 × 30 sec bursts, on ice). Protein concentration was measured using BCA protein assay kit (Thermo Fisher Scientific) and equal amounts of heavy and light lysates mixed in 1:1 ratio. Lysates were subsequently reduced with 5 mM dithiothreitol (DTT) for 30 min at room temperature and alkylated with 14 mM iodoacetamide for 30 min at room temperature in the dark. Following quenching with 5 mM DTT for 15 min in the dark lysates were diluted with 50 mM ammonium bicarbonate to reduce the concentration of urea to < 2M and digested with trypsin (Promega) overnight at 37°C. After digestion samples were acidified with trifluoroacetic acid (TFA) (Thermo Fisher Scientific) to a final concentration of 1 % (v/v), all insoluble material was removed by centrifugation and the supernatant was desalted with Sep-Pak C18 cartridges (Waters). The samples were further digested with LysC (Promega) for 2-3 h at 37°C and trypsin overnight at 37°C followed by desalting with Sep-Pak as above.

### Phosphopeptide enrichment

Desalted and vacuum dried samples were solubilized in 1 ml of loading buffer (80 % acetonitrile, 5 % TFA, 1 M glycolic acid) and mixed with 5 mg of TiO2 beads (Titansphere, 5 µm GL Sciences Japan). Samples were incubated for 10 min with agitation followed by a 1 min 2000 × g spin to pellet the beads. The supernatant containing all non-phosphorylated peptides (total proteome) was removed and stored at −80°C. The beads were washed with 150 μl of loading buffer followed by two additional wash steps, first with 150 μl 80 % acetonitrile, 1 % TFA and second with identical volume of 10 % acetonitrile, 0.2 % TFA. After each wash beads were pelleted by centrifugation (1 min at 2000 × g) and the supernatant discarded. The beads were dried in a vacuum centrifuge for 30 min followed by two elution steps at high pH. For the first elution step the beads were mixed with 100 μl of 1 % ammonium hydroxide (v/v) and for the second elution step with 100 µl of 5 % ammonium hydroxide (v/v). Each time the beads were incubated for 10 min with agitation and pelleted at 2000 × g for 1 min. The two elutions were combined and vacuum dried.

### Mass spectrometry sample fractionation and desalting

Both phospho- and total proteome (40 μg) samples were fractionated in a stage tip using Empore SDB-RPS discs (3M). Briefly, each stage tip was packed with one high performance extraction disc, samples were loaded in 100 μL of 1 % TFA, washed with 150 μL of 0.2 % TFA and eluted into 3 fractions with 100 μL of the following: 1) 100 mM ammonium formate, 20 % acetonitrile, 0.5 % formic acid; 2) 200 mM ammonium formate, 40 % acetonitrile, 0.5 % formic acid; 3) 5 % ammonium hydroxide, 60 % acetonitrile. The fractions were taken to dryness by vacuum centrifugation and further desalted on a stage tip using Empore C18 discs (3M). Briefly, each stage tip was packed with one C18 disc, conditioned with 100 µl of 100 % methanol, followed by 200 µl of 1 % TFA. The sample was loaded in 100 μL of 1 % TFA, washed 3 times with 200 µl of 1 % TFA and eluted with 50 µl of 50 % acetonitrile, 5 % TFA. The desalted peptides were vacuum dried in preparation for LC-MS/MS analysis.

### nLC-MS/MS and data processing

Samples were resuspended in 0.1 % TFA and loaded on a 50 cm Easy Spray PepMap column (75 μm inner diameter, 2 μm particle size, ThermoFisher Scientific) equipped with an integrated electrospray emitter. Reverse phase chromatography was performed using the RSLC nano U3000 (Thermo Fisher Scientific) with a binary buffer system (solvent A: 0.1% formic acid, 5% DMSO; solvent B: 80% acetonitrile, 0.1% formic acid, 5% DMSO) at a flow rate of 250 nl/min. The samples were run on a linear gradient of 2-35% B in 90 or 155 min with a total run time of 120 or 180 min, respectively, including column conditioning. The nanoLC was coupled to a Q Exactive mass spectrometer using an EasySpray nano source (Thermo Fisher Scientific). The Q Exactive was operated in data-dependent mode acquiring HCD MS/MS scans (R=17,500) after an MS1 scan (R=70,000) on the 10 most abundant ions using MS1 target of 1 × 106 ions, and MS2 target of 5 × 104 ions. The maximum ion injection time utilized for MS2 scans was 120 ms, the HCD normalized collision energy was set at 28, the dynamic exclusion was set at 20 or 30 s for 120 and 180 min runs, respectively, and the peptide match and isotope exclusion functions were enabled. Raw data files were processed with MaxQuant (82) (version 1.5.0.25) and peptides were identified from the MS/MS spectra searched against *Toxoplasma gondii* proteome (ToxoDB, 2017) using Andromeda (83) search engine. SILAC based experiments in MaxQuant were performed using the built-in quantification algorithm (82) with minimal ratio count = 1, enabled ‘Match between runs’ option for fractionated samples (time window 0.7 min) and ‘Re-quantify’ feature. Cysteine carbamidomethylation was selected as a fixed modification whereas methionine oxidation, acetylation of protein N-terminus and phosphorylation (S, T, Y) as variable modifications. The enzyme specificity was set to trypsin with maximum of 2 missed cleavages. The precursor mass tolerance was set to 20 ppm for the first search (used for mass re-calibration) and to 4.5 ppm for the main search. The datasets were filtered on posterior error probability to achieve 1% false discovery rate on protein, peptide and site level. “Unique and razor peptides” mode was selected to allow identification and quantification of proteins in groups (razor peptides are uniquely assigned to protein groups and not to individual proteins). Data were further analyzed as described in the Results section and in the Supplementary Table S4 using Microsoft Office Excel 2010 and Perseus (84) (version 1.5.0.9).

### Fractionation of PV membranes

Highly infected monolayers of HFFs were rinsed twice with phosphate buffered saline (PBS) and harvested. PBS containing 1 mM EDTA with protease inhibitors were added to the cells, and cells were mechanically disrupted by passage through a 27 g needle. Proteins secreted in the PV were separated by a low speed (2500 g) spin, and the resulting supernatant (LSS) was further separated by ultracentrifugation at 50,000 rpm for 2 hours at 4°C using TL100 rotor. The supernatant was aspirated as soluble fraction while the pellet was re-suspended in the same volume buffer. Equal volumes of each fraction were loaded on SDS-PAGE for analysis by western blot, which were quantified in ImageJ (85).

### Triton-X-114 partitioning

The LSS fraction of infected monolayers was prepared as above, and further partitioned using a protocol modified as follows from (54). Pre-condensed Triton-X-114 was added to the LSS to make the final concentration of 2% Triton-X-114. After a few minutes incubation on ice, the solution was warmed at 30°C for 3 minutes, then centrifuged for 5 minutes at 4000 rpm at room temperature. The top aqueous layer was collected in another tube, and added the same volume of 10 mM Tris-HCl pH7.4, 150 mM NaCl with protease inhibitors. The cleared solution after placed in 0°C for a few minutes was warmed at 30°C again, and centrifuged again. The aqueous layer was separated from the detergent enriched fraction. After separation, Triton-X-114 and buffer were added, respectively, to the aqueous and detergent phases in order to obtain equal volumes and approximately the same salt and surfactant content for both samples.

### Western blotting

Proteins were separated by SDS-PAGE and transferred to a PVDF membrane. Membranes were blocked for 1 hour in TBST + 3% milk, followed by overnight incubation at 4°C with primary antibody in blocking solution. The next day, membranes were washed 3× with TBST, followed by incubation at room temperature for 1-2 hours with HRP-conjugated secondary antibody (Sigma) in blocking buffer. After 3× washes with TBST, western blots were imaged using ECL Plus reagent (Pierce) on a GE ImageQuant LAS4000. Antibodies used in this study include: mouse anti-GRA1 (BioVision; 1:1,000 dilution), mouse anti-GRA2 (BioVision; 1:1,000 dilution), mouse anti-GRA3 (gift of J-F Dubremetz; 1:2,000 dilution), mouse anti-GRA4 (gift of LD Sibley; 1:10,000 dilution), mouse anti-GRA5 (BioVision; 1:1,000 dilution), mouse anti-GRA6 (gift of LD Sibley; 1:10,000 dilution), rabbit anti-GRA7 (gift of LD Sibley; 1:10,000 dilution), rabbit anti-ROP2 (1:10,000 dilution), rat anti-HA (Sigma; 1:500 dilution).

### Transmission electron microscopy

Cells were fixed on MatTek dishes with 2.5% (v/v) glutaraldehyde in 0.1M sodium cacodylate buffer. After three rinses in 0.1 M sodium cacodylate buffer, they were post-fixed with 1% osmium tetroxide and 0.8 % K_3_[Fe(CN_6_)] in 0.1 M sodium cacodylate buffer for 1 h at room temperature. Cells were rinsed with water and en bloc stained with 2% aqueous uranyl acetate overnight. After three rinses with water, specimens were dehydrated with increasing concentration of ethanol, infiltrated with Embed-812 resin and polymerized in a 70ºC oven overnight. Blocks were sectioned with a diamond knife (Diatome) on a Leica Ultracut UC7 ultramicrotome (Leica Microsystems) and collected onto copper grids, post stained with 2% Uranyl acetate in water and lead citrate. Images were acquired on a Tecnai G2 spirit transmission electron microscope (FEI) equipped with a LaB_6_ source at 120 kV. Images were analyzed and quantified using the Fiji distribution of ImageJ (85).

### Figure generation

Structural models were generated using PyMOL v1.7 (86). Secondary structure cartoons in Figure 3 were generated using the Pro-origami web server (87). Data plotting and statistical analyses were conducted in Graphpad Prism v7.02. All figures were created in Inkscape v0.91.

## Acknowledgments

MLR acknowledges funding from the Welch Foundation (I-1936-20170325) and NSF (MCB1553334). TB is funded, in part, by NIH training grant T32GM008203. MT was supported by awards of the United States NIH (NIH-R01AI123457) and The Francis Crick Institute (https://www.crick.ac.uk/), which receives its core funding from Cancer Research UK (FC001189; https://www.cancerresearchuk.org), the UK Medical Research Council (FC001189; https://www.mrc.ac.uk/) and the Wellcome Trust (FC001189; https://wellcome.ac.uk/). DB acknowledges funding from the NIH (R01GM117080 and R21GM126406). We thank V. Tagliabracci and V. Muralidharan for helpful comments on the manuscript.

---

**Figure S1a:**
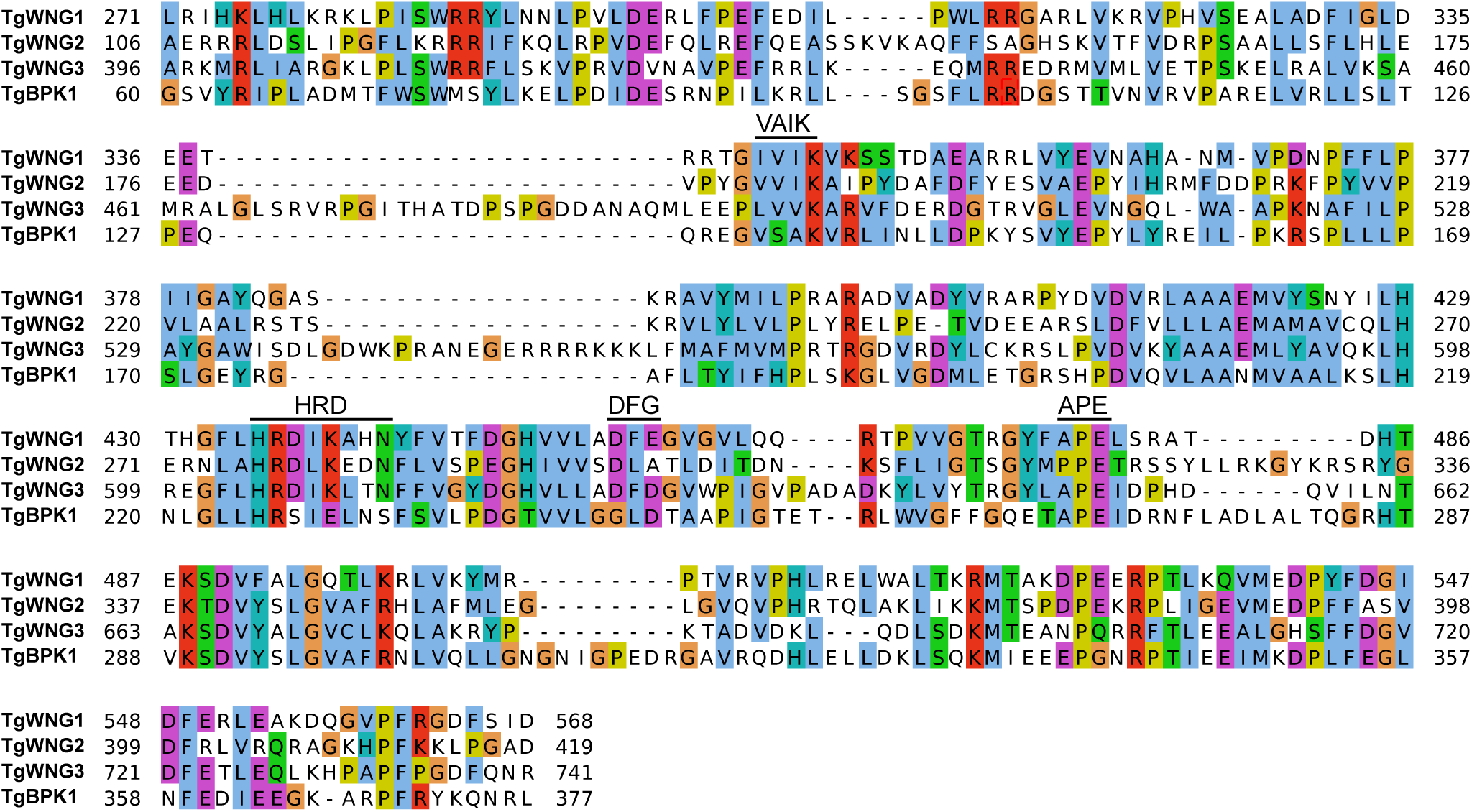
Alignment of *Toxoplasma gondii* WNG kinase domains. Canonical kinase motifs are indicated above sequences.

**Figure S1b:**
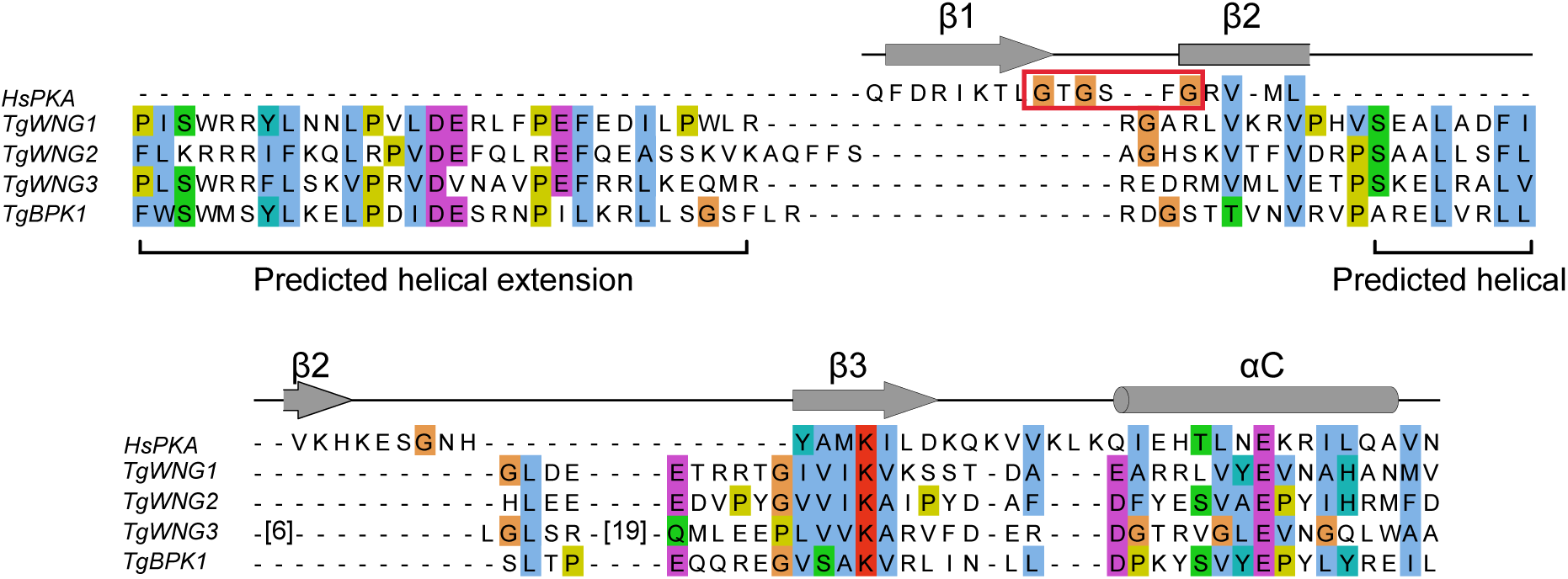
Alignment of the N-lobes of human PKA and *Toxoplasma gondii* WNG kinase domains. Secondary structure elements from the PKA crystal structure (1ATP) are indicated as cartoons above the alignment. The PKA Gly-loop is boxed in red. Note that the WNG kinases lack sequences corresponding to the Gly-loop, which has been replaced with a conserved sequence predicted to form helices.

**Figure S3:**
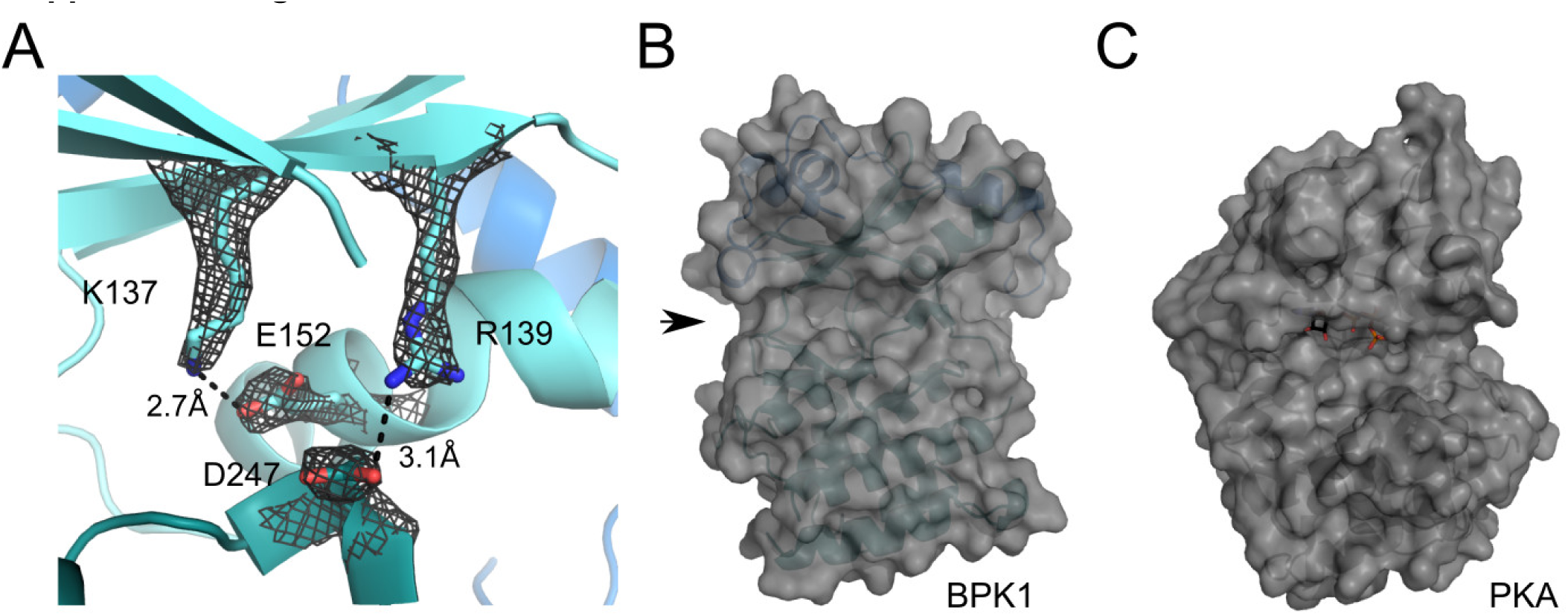
BPK1 has a divergent, open active site. (A) BPK1 active site superposed with the 2*FO– FC* electron density map contoured at 2σ. Two salt bridges are highlighted as sticks: the conserved bridge between the αC E152 and the VAIK K137 as well as an unusual, WNG family-specific salt bridge between R139 and D247 (an acidic substitution at the DFG Gly position). The lack of Gly-loop creates an open active site in BPK1, indicated with an arrow in (B). This is compared to the more restricted active site in canonical kinases such as PKA, shown in (C). Note that the two kinases are shown in equivalent orientations.

**Figure S4:**
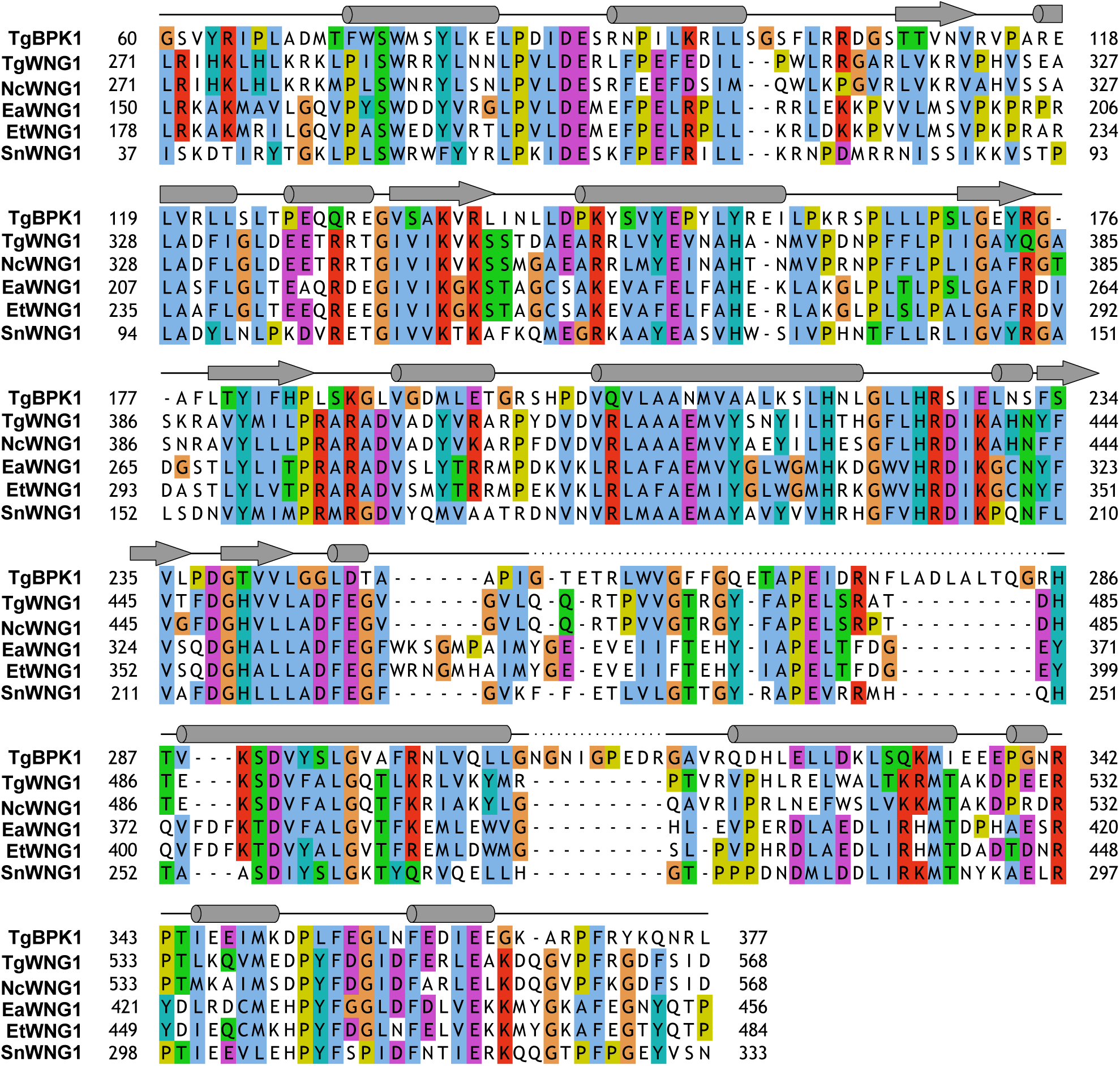
Multiple sequence alignment of WNG1 kinase domains with TgBPK1. The secondary structure from the TgBPK1 crystal structure are shown in cartoon above the alignment. Dashed lines indicate a lack of density corresponding to the indicated sequence.

**Figure S5:**
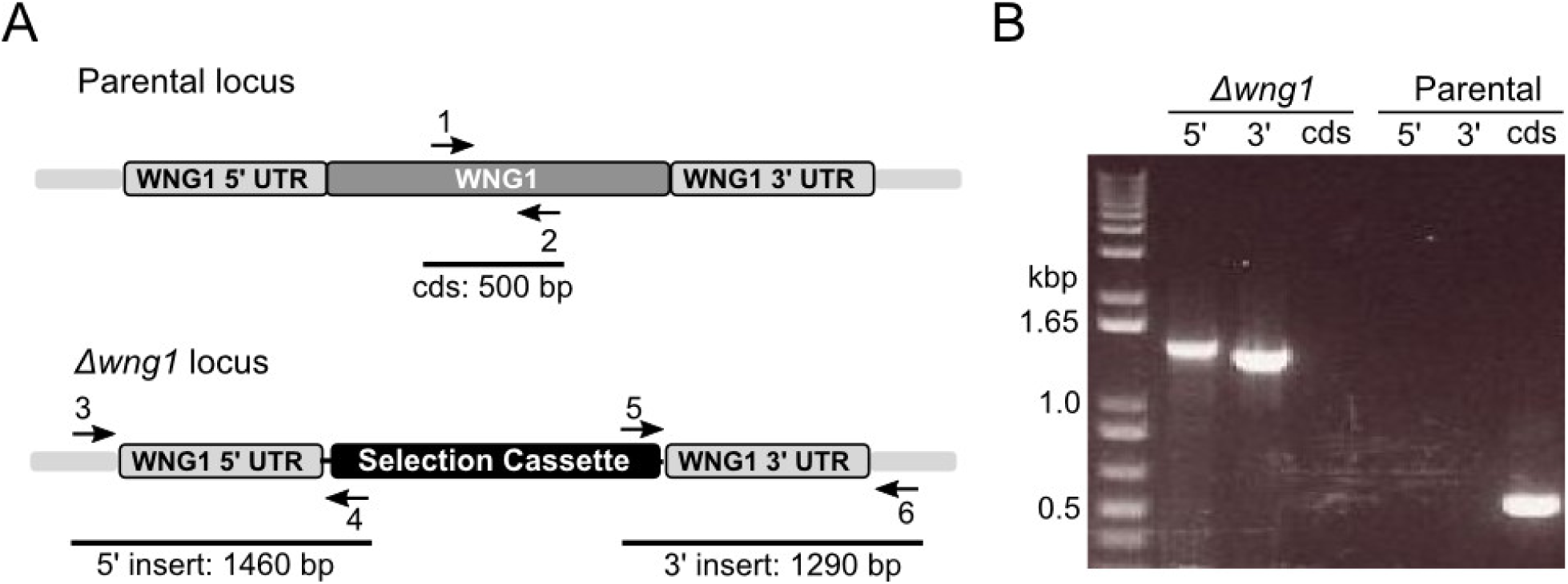
Generation of WNG1 knockout parasites. RH*Δwng1* parasites were generated by double homologous recombination in which the WNG1 genomic sequence was replaced by a HXGPRT selection cassette. (A) Cartoon of parental and knockout loci indicating binding sites for primers used to verify knockout. (B) PCR demonstrating insertion of selection cassette and loss of coding sequence (cds) in knockout parasites. Primers sequences are listed in Supplemental Table S10.

**Figure S5:**
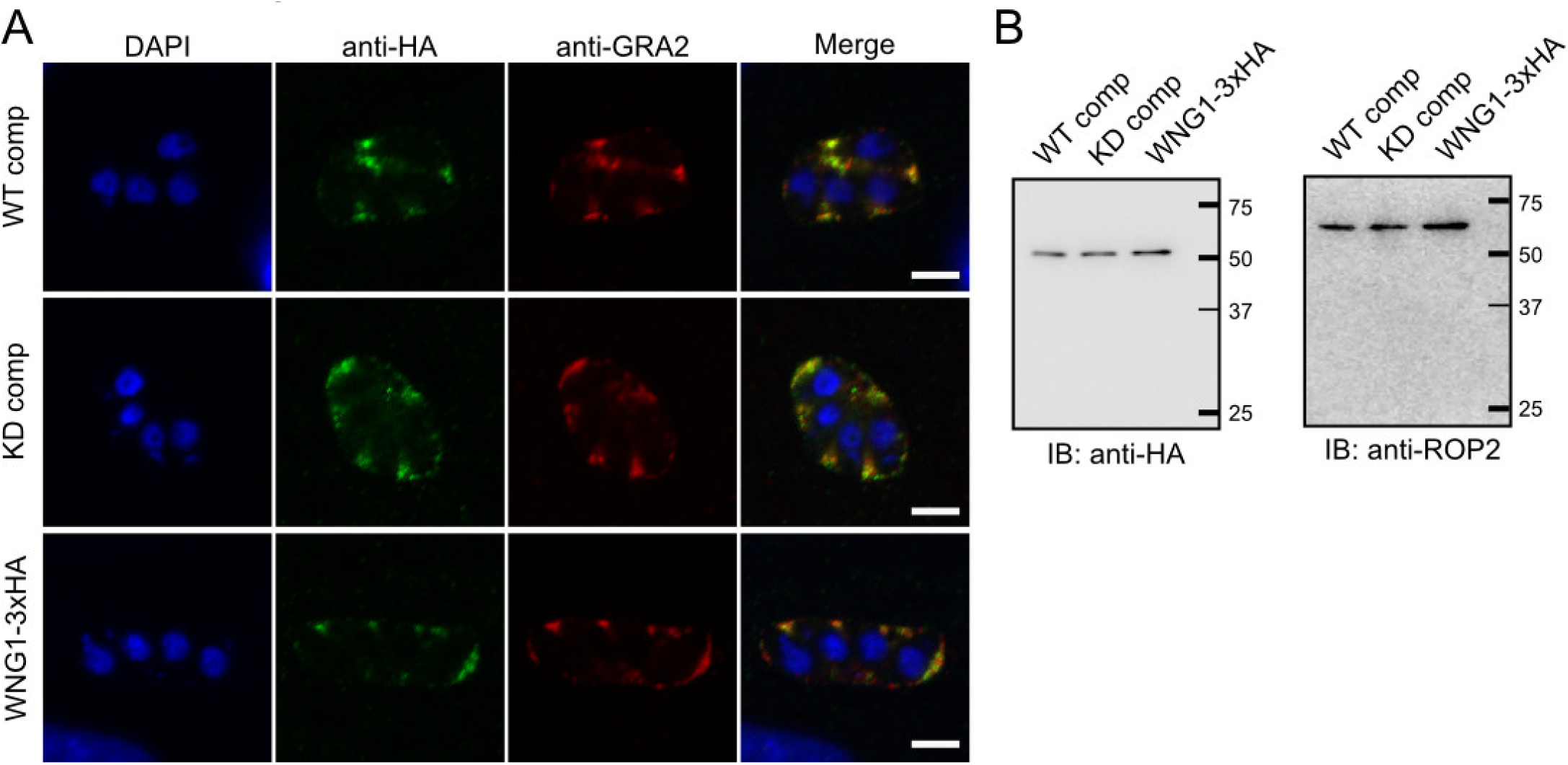
WNG1 complements faithfully localize to dense granules and PV. (A) 0.5 μm confocal slices of the wild-type (WT) and kinase-dead (KD) WNG1 complemented parasites as well as the endogenously tagged WNG1-3xHA were stained with DAPI (blue), anti-HA (green), and the dense granule and IVN marker GRA2 (red). (B) Both the WT and KD WNG1-complements are expressed at similar levels to the endogenously 3xHA tagged protein, as demonstrated by western blot, using ROP2 as a loading control.

**Figure S5:**
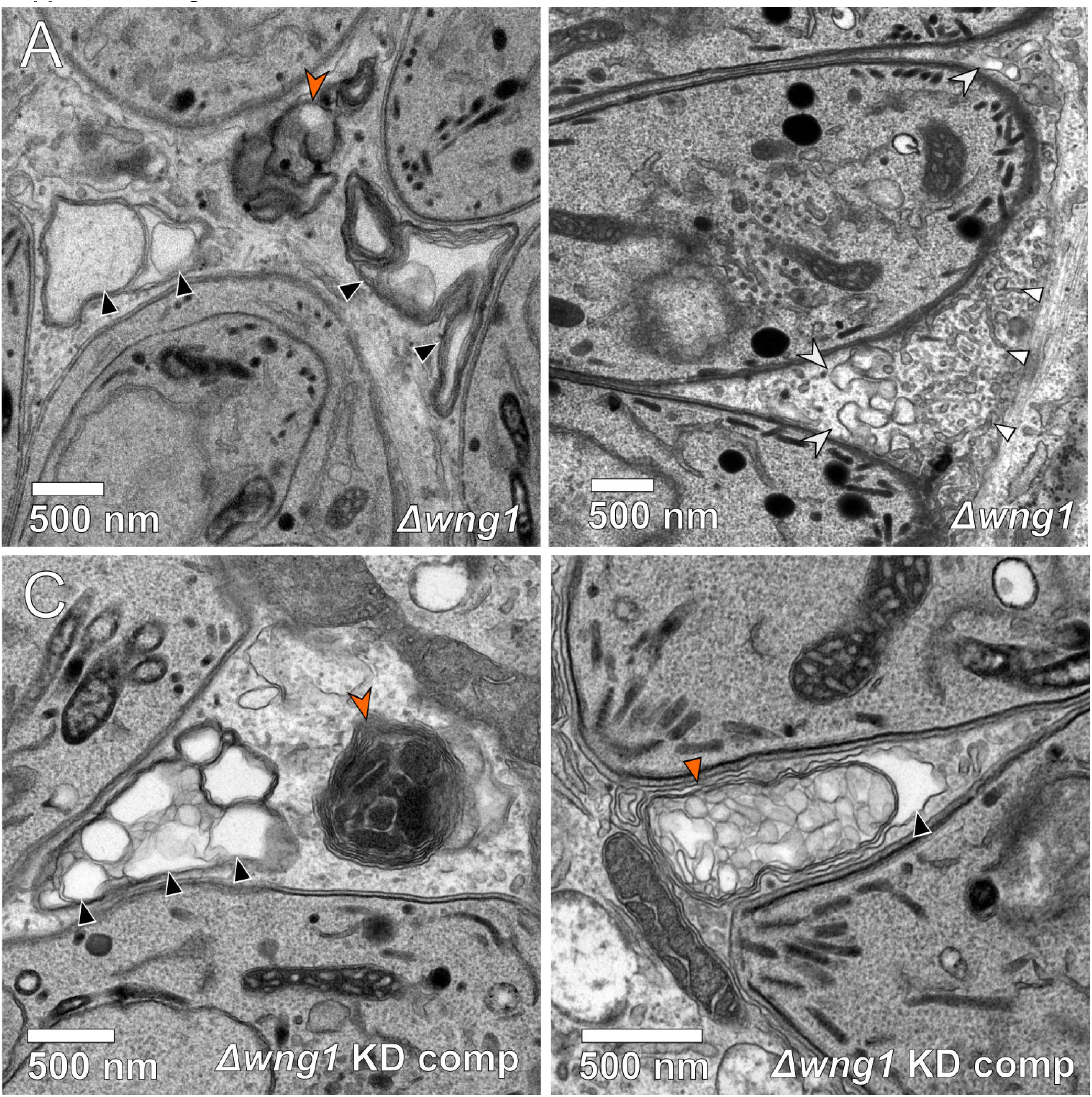
Unusual membrane structures in vacuoles lacking active WNG1 kinase. Representative transmission electron microscopic images of (A,B) RH*Δwng1* and (C,D) RH*Δwng1* complemented with kinase-dead WNG1. IVN tubules are indicated with small white triangles. Multilamellar vesicles are indicated with small solid orange triangles. Multilamellar structures in which internal vesicles appear to have been lost during fixation or to have collapsed into sheets are indicated with black triangles. Electron dense multilamellar structure are indicated with a large orange arrowheads in (A) and (C). Membrane “whirls” that appear connected with IVN tubules are indicated with large white arrowheads in (B).

**Figure S8:**
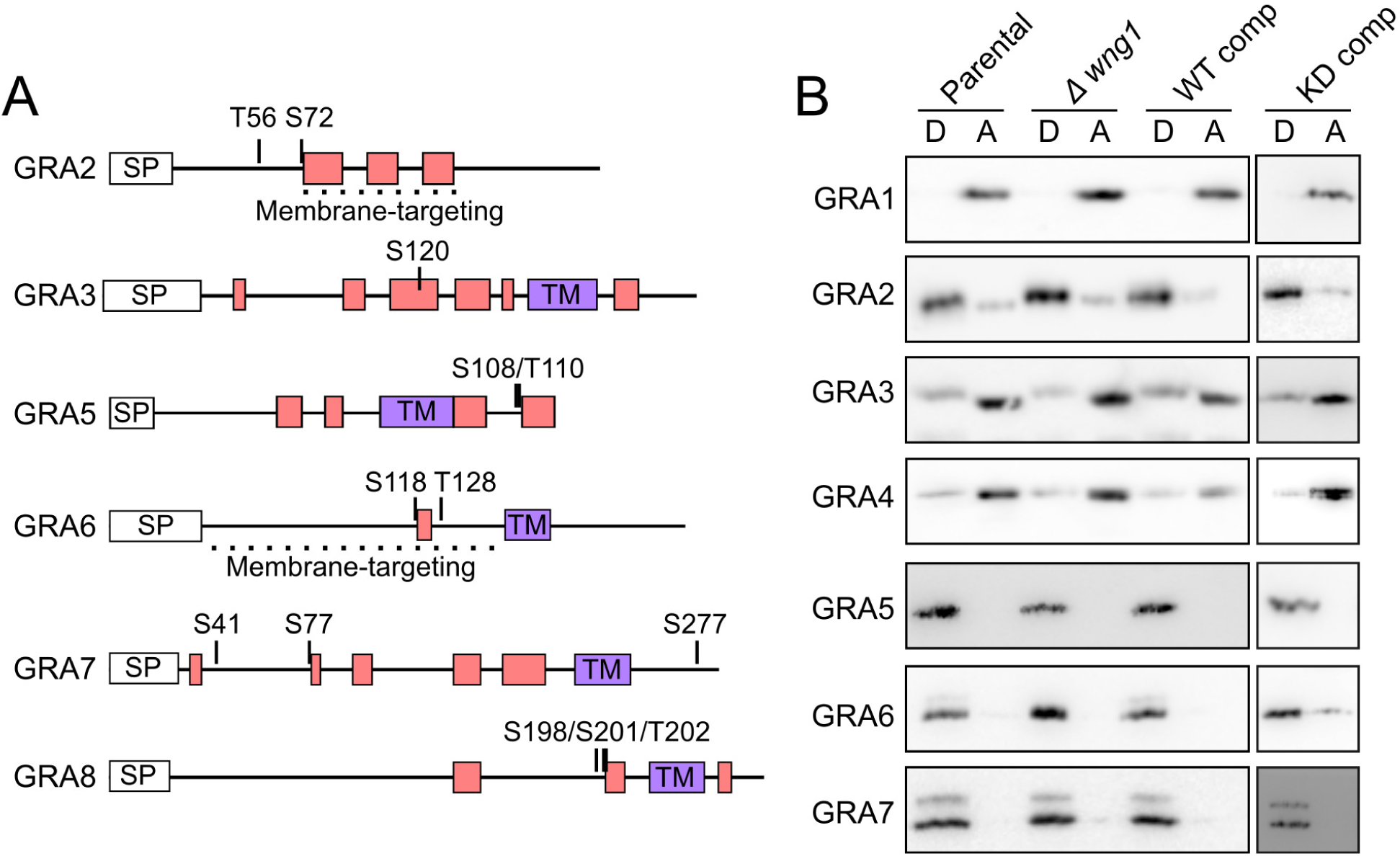
WNG1 activity does not affect TX-114 partitioning of IVN GRA proteins. (A) WNG1-dependent phosphosites are mapped onto the predicted secondary structures of the indicated GRA proteins. Predicted α-helices are shown as rectangles. Predicted transmembrane helices (TM) are shaded purple. (B) Host and PV membranes from cells infected with the indicated strains were partitioned in TX-114 and the detergent (D) and aqueous (A) phases were separated by SDS-PAGE and analyzed by western blot probed with antibodies to the indicated GRA proteins.

**Table S1:**
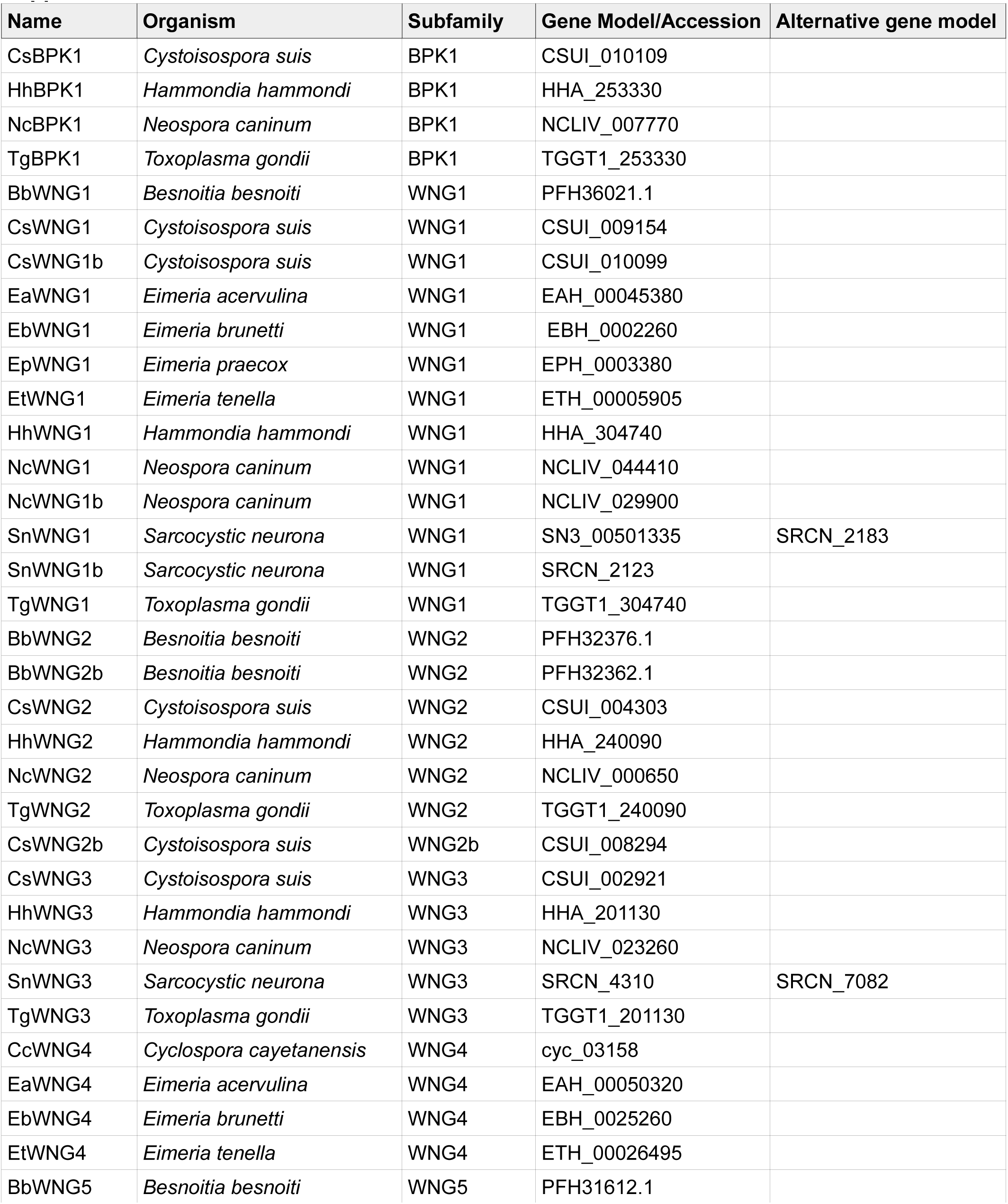
Gene models (for sequences in ToxoDB) or NCBI accession numbers of sequences used in this study.

## References

1. Manning G, Whyte DB, Martinez R, Hunter T, Sudarsanam S. 2002. The Protein Kinase Complement of the Human Genome. Science 298:1912–1934.

2. Pappas G, Roussos N, Falagas ME. 2009. Toxoplasmosis snapshots: global status of Toxoplasma gondii seroprevalence and implications for pregnancy and congenital toxoplasmosis. Int J Parasitol 39:1385–1394.

3. Boothroyd JC, Dubremetz J-F. 2008. Kiss and spit: the dual roles of Toxoplasma rhoptries. Nat Rev Microbiol 6:79–88.

4. Hakimi M-A, Olias P, Sibley LD. 2017. Toxoplasma Effectors Targeting Host Signaling and Transcription. Clin Microbiol Rev 30:615–645.

5. Peixoto L, Chen F, Harb OS, Davis PH, Beiting DP, Brownback CS, Ouloguem D, Roos DS. 2010. Integrative genomic approaches highlight a family of parasite-specific kinases that regulate host responses. Cell Host Microbe 8:208–218.

6. Boothroyd JC. 2013. Have it your way: how polymorphic, injected kinases and pseudokinases enable Toxoplasma to subvert host defenses. PLoS Pathog 9:e1003296.

7. Mordue DG, Håkansson S, Niesman I, Sibley LD. 1999. Toxoplasma gondii resides in a vacuole that avoids fusion with host cell endocytic and exocytic vesicular trafficking pathways. Exp Parasitol 92:87–99.

8. Hunn JP, Feng CG, Sher A, Howard JC. 2011. The immunity-related GTPases in mammals: a fast-evolving cell-autonomous resistance system against intracellular pathogens. Mamm Genome Off J Int Mamm Genome Soc 22:43–54.

9. Clough B, Wright JD, Pereira PM, Hirst EM, Johnston AC, Henriques R, Frickel E-M. 2016. K63-Linked Ubiquitination Targets Toxoplasma gondii for Endo-lysosomal Destruction in IFNγ-Stimulated Human Cells. PLoS Pathog 12:e1006027.

10. Etheridge RD, Alaganan A, Tang K, Lou HJ, Turk BE, Sibley LD. 2014. The Toxoplasma pseudokinase ROP5 forms complexes with ROP18 and ROP17 kinases that synergize to control acute virulence in mice. Cell Host Microbe 15:537–550.

11. Reese ML, Shah N, Boothroyd JC. 2014. The Toxoplasma Pseudokinase ROP5 Is an Allosteric Inhibitor of the Immunity-related GTPases. J Biol Chem 289:27849–27858.

12. Franco M, Panas MW, Marino ND, Lee M-CW, Buchholz KR, Kelly FD, Bednarski JJ, Sleckman BP, Pourmand N, Boothroyd JC. 2016. A Novel Secreted Protein, MYR1, Is Central to Toxoplasma’s Manipulation of Host Cells. mBio 7:e02231–15.

13. Schwab JC, Beckers CJ, Joiner KA. 1994. The parasitophorous vacuole membrane surrounding intracellular Toxoplasma gondii functions as a molecular sieve. Proc Natl Acad Sci U S A 91:509–513.

14. Gold DA, Kaplan AD, Lis A, Bett GCL, Rosowski EE, Cirelli KM, Bougdour A, Sidik SM, Beck JR, Lourido S, Egea PF, Bradley PJ, Hakimi M-A, Rasmusson RL, Saeij JPJ. 2015. The Toxoplasma Dense Granule Proteins GRA17 and GRA23 Mediate the Movement of Small Molecules between the Host and the Parasitophorous Vacuole. Cell Host Microbe 17:642–652.

15. Sibley LD, Niesman IR, Parmley SF, Cesbron-Delauw MF. 1995. Regulated secretion of multi-lamellar vesicles leads to formation of a tubulo-vesicular network in host-cell vacuoles occupied by Toxoplasma gondii. J Cell Sci 108 (Pt 4):1669–1677.

16. Coppens I, Dunn JD, Romano JD, Pypaert M, Zhang H, Boothroyd JC, Joiner KA. 2006. Toxoplasma gondii sequesters lysosomes from mammalian hosts in the vacuolar space. Cell 125:261–274.

17. Romano JD, Sonda S, Bergbower E, Smith ME, Coppens I. 2013. Toxoplasma gondii salvages sphingolipids from the host Golgi through the rerouting of selected Rab vesicles to the parasitophorous vacuole. Mol Biol Cell 24:1974–1995.

18. Dou Z, McGovern OL, Cristina MD, Carruthers VB. 2014. Toxoplasma gondii Ingests and Digests Host Cytosolic Proteins. mBio5:e01188–14.

19. Lopez J, Bittame A, Massera C, Vasseur V, Effantin G, Valat A, Buaillon C, Allart S, Fox BA, Rommereim LM, Bzik DJ, Schoehn G, Weissenhorn W, Dubremetz J-F, Gagnon J, Mercier C, Cesbron-Delauw M-F, Blanchard N. 2015. Intravacuolar Membranes Regulate CD8 T Cell Recognition of Membrane-Bound Toxoplasma gondii Protective Antigen. Cell Rep 13:2273–2286.

20. Reese ML, Boothroyd JC. 2009. A helical membrane-binding domain targets the Toxoplasma ROP2 family to the parasitophorous vacuole. Traffic 10:1458–1470.

21. Fentress SJ, Steinfeldt T, Howard JC, Sibley LD. 2012. The arginine-rich N-terminal domain of ROP18 is necessary for vacuole targeting and virulence of Toxoplasma gondii. Cell Microbiol 14:1921–1933.

22. Mercier C, Howe DK, Mordue D, Lingnau M, Sibley LD. 1998. Targeted disruption of the GRA2 locus in Toxoplasma gondii decreases acute virulence in mice. Infect Immun 66:4176–4182.

23. Treeck M, Sanders JL, Elias JE, Boothroyd JC. 2011. The phosphoproteomes of Plasmodium falciparum and Toxoplasma gondii reveal unusual adaptations within and beyond the parasites’ boundaries. Cell Host Microbe 10:410–419.

24. Saeij JPJ, Coller S, Boyle JP, Jerome ME, White MW, Boothroyd JC. 2007. Toxoplasma co-opts host gene expression by injection of a polymorphic kinase homologue. Nature 445:324–327.

25. Reese ML, Zeiner GM, Saeij JPJ, Boothroyd JC, Boyle JP. 2011. Polymorphic family of injected pseudokinases is paramount in Toxoplasma virulence. Proc Natl Acad Sci U S A 108:9625–9630.

26. Bradley PJ, Sibley LD. 2007. Rhoptries: an arsenal of secreted virulence factors. Curr Opin Microbiol 10:582–587.

27. Jones NG, Wang Q, Sibley LD. 2016. Secreted Protein Kinases Regulate Cyst Burden During Chronic Toxoplasmosis. Cell Microbiol.

28. Walker JE, Saraste M, Runswick MJ, Gay NJ. 1982. Distantly related sequences in the alpha- and beta-subunits of ATP synthase, myosin, kinases and other ATP-requiring enzymes and a common nucleotide binding fold. EMBO J 1:945–951.

29. Hanks SK, Hunter T. 1995. Protein kinases 6. The eukaryotic protein kinase superfamily: kinase (catalytic) domain structure and classification. FASEB J Off Publ Fed Am Soc Exp Biol 9:576–596.

30. Buchholz KR, Bowyer PW, Boothroyd JC. 2013. Bradyzoite pseudokinase 1 is crucial for efficient oral infectivity of the Toxoplasma gondii tissue cyst. Eukaryot Cell 12:399–410.

31. Talevich E, Kannan N. 2013. Structural and evolutionary adaptation of rhoptry kinases and pseudokinases, a family of coccidian virulence factors. BMC Evol Biol 13:117.

32. Fox BA, Sanders KL, Rommereim LM, Guevara RB, Bzik DJ. 2016. Secretion of Rhoptry and Dense Granule Effector Proteins by Nonreplicating Toxoplasma gondii Uracil Auxotrophs Controls the Development of Antitumor Immunity. PLoS Genet 12:e1006189.

33. Ossorio PN, Schwartzman JD, Boothroyd JC. 1992. A Toxoplasma gondii rhoptry protein associated with host cell penetration has unusual charge asymmetry. Mol Biochem Parasitol 50:1–15.

34. Brennan DF, Dar AC, Hertz NT, Chao WCH, Burlingame AL, Shokat KM, Barford D. 2011. A Raf-induced allosteric transition of KSR stimulates phosphorylation of MEK. Nature 472:366–369.

35. Murphy JM, Czabotar PE, Hildebrand JM, Lucet IS, Zhang J-G, Alvarez-Diaz S, Lewis R, Lalaoui N, Metcalf D, Webb AI, Young SN, Varghese LN, Tannahill GM, Hatchell EC, Majewski IJ, Okamoto T, Dobson RCJ, Hilton DJ, Babon JJ, Nicola NA, Strasser A, Silke J, Alexander WS. 2013. The pseudokinase MLKL mediates necroptosis via a molecular switch mechanism. Immunity 39:443–453.

36. Zhang H, Zhu Q, Cui J, Wang Y, Chen MJ, Guo X, Tagliabracci VS, Dixon JE, Xiao J. 2018. Structure and evolution of the Fam20 kinases. Nat Commun 9:1218.

37. Zhu Q, Venzke D, Walimbe AS, Anderson ME, Fu Q, Kinch LN, Wang W, Chen X, Grishin NV, Huang N, Yu L, Dixon JE, Campbell KP, Xiao J. 2016. Structure of protein O-mannose kinase reveals a unique active site architecture. eLife 5.

38. Murphy JM, Zhang Q, Young SN, Reese ML, Bailey FP, Eyers PA, Ungureanu D, Hammaren H, Silvennoinen O, Varghese LN, Chen K, Tripaydonis A, Jura N, Fukuda K, Qin J, Nimchuk Z, Mudgett MB, Elowe S, Gee CL, Liu L, Daly RJ, Manning G, Babon JJ, Lucet IS. 2014. A robust methodology to subclassify pseudokinases based on their nucleotide-binding properties. Biochem J 457:323–334.

39. Levinson NM, Kuchment O, Shen K, Young MA, Koldobskiy M, Karplus M, Cole PA, Kuriyan J. 2006. A Src-like inactive conformation in the abl tyrosine kinase domain. PLoS Biol 4:e144.

40. Ruff EF, Muretta JM, Thompson AR, Lake EW, Cyphers S, Albanese SK, Hanson SM, Behr JM, Thomas DD, Chodera JD, Levinson NM. 2018. A dynamic mechanism for allosteric activation of Aurora kinase A by activation loop phosphorylation. eLife 7.

41. Huang H, Zeqiraj E, Dong B, Jha BK, Duffy NM, Orlicky S, Thevakumaran N, Talukdar M, Pillon MC, Ceccarelli DF, Wan LCK, Juang Y-C, Mao DYL, Gaughan C, Brinton MA, Perelygin AA, Kourinov I, Guarné A, Silverman RH, Sicheri F. 2014. Dimeric structure of pseudokinase RNase L bound to 2-5A reveals a basis for interferon-induced antiviral activity. Mol Cell 53:221–234.

42. Rudolf AF, Skovgaard T, Knapp S, Jensen LJ, Berthelsen J. 2014. A comparison of protein kinases inhibitor screening methods using both enzymatic activity and binding affinity determination. PloS One 9:e98800.

43. Yoshida T, Kakizuka A, Imamura H. 2016. BTeam, a Novel BRET-based Biosensor for the Accurate Quantification of ATP Concentration within Living Cells. Sci Rep 6:39618.

44. Fox BA, Rommereim LM, Guevara RB, Falla A, Hortua Triana MA, Sun Y, Bzik DJ. 2016. The Toxoplasma gondii Rhoptry Kinome Is Essential for Chronic Infection. mBio 7.

45. Mercier C, Dubremetz J-F, Rauscher B, Lecordier L, Sibley LD, Cesbron-Delauw M-F. 2002. Biogenesis of nanotubular network in Toxoplasma parasitophorous vacuole induced by parasite proteins. Mol Biol Cell 13:2397–2409.

46. Treeck M, Sanders JL, Gaji RY, LaFavers KA, Child MA, Arrizabalaga G, Elias JE, Boothroyd JC. 2014. The calcium-dependent protein kinase 3 of toxoplasma influences basal calcium levels and functions beyond egress as revealed by quantitative phosphoproteome analysis. PLoS Pathog 10:e1004197.

47. Nadipuram SM, Kim EW, Vashisht AA, Lin AH, Bell HN, Coppens I, Wohlschlegel JA, Bradley PJ. 2016. In Vivo Biotinylation of the Toxoplasma Parasitophorous Vacuole Reveals Novel Dense Granule Proteins Important for Parasite Growth and Pathogenesis. mBio 7.

48. Labruyere E, Lingnau M, Mercier C, Sibley LD. 1999. Differential membrane targeting of the secretory proteins GRA4 and GRA6 within the parasitophorous vacuole formed by Toxoplasma gondii. Mol Biochem Parasitol 102:311–324.

49. Lecordier L, Mercier C, Sibley LD, Cesbron-Delauw MF. 1999. Transmembrane insertion of the Toxoplasma gondii GRA5 protein occurs after soluble secretion into the host cell. Mol Biol Cell 10:1277–1287.

50. Mercier C, Cesbron-Delauw MF, Sibley LD. 1998. The amphipathic alpha helices of the toxoplasma protein GRA2 mediate post-secretory membrane association. J Cell Sci 111 (Pt 15):2171–2180.

51. Jacobs D, Dubremetz JF, Loyens A, Bosman F, Saman E. 1998. Identification and heterologous expression of a new dense granule protein (GRA7) from Toxoplasma gondii. Mol Biochem Parasitol 91:237–249.

52. Carey KL, Donahue CG, Ward GE. 2000. Identification and molecular characterization of GRA8, a novel, proline-rich, dense granule protein of Toxoplasma gondii. Mol Biochem Parasitol 105:25–37.

53. Ossorio PN, Dubremetz JF, Joiner KA. 1994. A soluble secretory protein of the intracellular parasite Toxoplasma gondii associates with the parasitophorous vacuole membrane through hydrophobic interactions. J Biol Chem 269:15350–15357.

54. Bordier C. 1981. Phase separation of integral membrane proteins in Triton X-114 solution. J Biol Chem 256:1604–1607.

55. Gendrin C, Bittame A, Mercier C, Cesbron-Delauw M-F. 2010. Post-translational membrane sorting of the Toxoplasma gondii GRA6 protein into the parasite-containing vacuole is driven by its N-terminal domain. Int J Parasitol 40:1325–1334.

56. Travier L, Mondragon R, Dubremetz J-F, Musset K, Mondragon M, Gonzalez S, Cesbron-Delauw M-F, Mercier C. 2008. Functional domains of the Toxoplasma GRA2 protein in the formation of the membranous nanotubular network of the parasitophorous vacuole. Int J Parasitol 38:757–773.

57. Middelbeek J, Clark K, Venselaar H, Huynen MA, van Leeuwen FN. 2010. The alpha-kinase family: an exceptional branch on the protein kinase tree. Cell Mol Life Sci CMLS 67:875–890.

58. Vaillancourt JP, Lyons C, Côté GP. 1988. Identification of two phosphorylated threonines in the tail region of Dictyostelium myosin II. J Biol Chem 263:10082–10087.

59. Yamaguchi H, Matsushita M, Nairn AC, Kuriyan J. 2001. Crystal structure of the atypical protein kinase domain of a TRP channel with phosphotransferase activity. Mol Cell 7:1047–1057.

60. Drennan D, Ryazanov AG. 2004. Alpha-kinases: analysis of the family and comparison with conventional protein kinases. Prog Biophys Mol Biol 85:1–32.

61. Lee S-N, Lindberg I. 2008. 7B2 prevents unfolding and aggregation of prohormone convertase 2. Endocrinology 149:4116–4127.

62. Lee S-N, Hwang JR, Lindberg I. 2006. Neuroendocrine protein 7B2 can be inactivated by phosphorylation within the secretory pathway. J Biol Chem 281:3312–3320.

63. Ramos-Molina B, Lindberg I. 2015. Phosphorylation and Alternative Splicing of 7B2 Reduce Prohormone Convertase 2 Activation. Mol Endocrinol Baltim Md 29:756–764.

64. Craver MPJ, Knoll LJ. 2007. Increased efficiency of homologous recombination in Toxoplasma gondii dense granule protein 3 demonstrates that GRA3 is not necessary in cell culture but does contribute to virulence. Mol Biochem Parasitol 153:149–157.

65. Camacho C, Coulouris G, Avagyan V, Ma N, Papadopoulos J, Bealer K, Madden TL. 2009. BLAST+: architecture and applications. BMC Bioinformatics 10:421.

66. Katoh K, Standley DM. 2013. MAFFT multiple sequence alignment software version 7: improvements in performance and usability. Mol Biol Evol 30:772–780.

67. Stamatakis A. 2014. RAxML version 8: a tool for phylogenetic analysis and post-analysis of large phylogenies. Bioinforma Oxf Engl 30:1312–1313.

68. Smits SA, Ouverney CC. 2010. jsPhyloSVG: a javascript library for visualizing interactive and vector-based phylogenetic trees on the web. PloS One 5:e12267.

69. Huynh M, Carruthers VB. 2009. Tagging of endogenous genes in a Toxoplasma gondii strain lacking Ku80. Eukaryot Cell 8:530–9.

70. Donald RG, Carter D, Ullman B, Roos DS. 1996. Insertional tagging, cloning, and expression of the Toxoplasma gondii hypoxanthine-xanthine-guanine phosphoribosyltransferase gene. Use as a selectable marker for stable transformation. J Biol Chem 271:14010–14019.

71. Soldati D, Kim K, Kampmeier J, Dubremetz JF, Boothroyd JC. 1995. Complementation of a Toxoplasma gondii ROP1 knock-out mutant using phleomycin selection. Mol Biochem Parasitol 74:87–97.

72. Otwinowski Z, Minor W. 1997. Processing of X-ray diffraction data collected in oscillation mode. Methods Enzymol 276:307–326.

73. Sheldrick GM. 2008. A short history of SHELX. Acta Crystallogr A 64:112–122.

74. Terwilliger TC, Berendzen J. 1999. Automated MAD and MIR structure solution. Acta Crystallogr D Biol Crystallogr 55:849–861.

75. Terwilliger T. 2004. SOLVE and RESOLVE: automated structure solution, density modification and model building. J Synchrotron Radiat 11:49–52.

76. Adams PD, Afonine PV, Bunkóczi G, Chen VB, Davis IW, Echols N, Headd JJ, Hung L-W, Kapral GJ, Grosse-Kunstleve RW, McCoy AJ, Moriarty NW, Oeffner R, Read RJ, Richardson DC, Richardson JS, Terwilliger TC, Zwart PH. 2010. PHENIX: a comprehensive Python-based system for macromolecular structure solution. Acta Crystallogr D Biol Crystallogr 66:213–221.

77. Emsley P, Lohkamp B, Scott WG, Cowtan K. 2010. Features and development of Coot. Acta Crystallogr D Biol Crystallogr 66:486–501.

78. Winn MD, Murshudov GN, Papiz MZ. 2003. Macromolecular TLS refinement in REFMAC at moderate resolutions. Methods Enzymol 374:300–321.

79. Chen VB, Arendall WB, Headd JJ, Keedy DA, Immormino RM, Kapral GJ, Murray LW, Richardson JS, Richardson DC. 2009. *MolProbity*?: all-atom structure validation for macromolecular crystallography. Acta Crystallogr D Biol Crystallogr 66:12–21.

80. Sali A, Blundell TL. 1993. Comparative protein modelling by satisfaction of spatial restraints. J Mol Biol 234:779–815.

81. Sievers F, Wilm A, Dineen D, Gibson TJ, Karplus K, Li W, Lopez R, McWilliam H, Remmert M, Söding J, Thompson JD, Higgins DG. 2011. Fast, scalable generation of high-quality protein multiple sequence alignments using Clustal Omega. Mol Syst Biol 7:539.

82. Cox J, Mann M. 2008. MaxQuant enables high peptide identification rates, individualized p.p.b.- range mass accuracies and proteome-wide protein quantification. Nat Biotechnol 26:1367–1372.

83. Cox J, Neuhauser N, Michalski A, Scheltema RA, Olsen JV, Mann M. 2011. Andromeda: a peptide search engine integrated into the MaxQuant environment. J Proteome Res 10:1794–1805.

84. Tyanova S, Temu T, Sinitcyn P, Carlson A, Hein MY, Geiger T, Mann M, Cox J. 2016. The Perseus computational platform for comprehensive analysis of (prote)omics data. Nat Methods 13:731–740.

85. Schindelin J, Arganda-Carreras I, Frise E, Kaynig V, Longair M, Pietzsch T, Preibisch S, Rueden C, Saalfeld S, Schmid B, Tinevez J-Y, White DJ, Hartenstein V, Eliceiri K, Tomancak P, Cardona A. 2012. Fiji: an open-source platform for biological-image analysis. Nat Methods 9:676–682.

86. Schrödinger, LLC. 2015. The PyMOL Molecular Graphics System, Version 1.7.

87. Stivala A, Wybrow M, Wirth A, Whisstock JC, Stuckey PJ. 2011. Automatic generation of protein structure cartoons with Pro-origami. Bioinforma Oxf Engl 27:3315–3316.

